# A framework for evaluating the performance of SMLM cluster analysis algorithms

**DOI:** 10.1101/2021.06.19.449098

**Authors:** Daniel J. Nieves, Jeremy A. Pike, Florian Levet, Juliette Griffié, Daniel Sage, Edward A.K. Cohen, Jean-Baptiste Sibarita, Mike Heilemann, Dylan M. Owen

## Abstract

Single molecule localisation microscopy (SMLM) generates data in the form of Cartesian coordinates of localised fluorophores. Cluster analysis is an attractive route for extracting biologically meaningful information from such data and has been widely applied. Despite the range of developed cluster analysis algorithms, there exists no consensus framework for the evaluation of their performance. Here, we use a systematic approach based on two metrics, the Adjusted Rand Index (ARI) and Intersection over Union (IoU), to score the success of clustering algorithms in diverse simulated clustering scenarios mimicking experimental data. We demonstrate the framework using three analysis algorithms: DBSCAN, ToMATo and KDE, show how to deduce optimal analysis parameters and how they are affected by fluorophore multiple blinking. We propose that these standard conditions and metrics become the basis for future analysis algorithm development and evaluation.

## Introduction

Single molecule localisation microscopy (SMLM) is now an established and widely used technique^1^. The assembly of biomolecules into aggregates (clusters) is a key process in cell biology^2,3^, and SMLM provides a route to study these complexes^1^. Different SMLM modalities can be distinguished by the way the sparse pointillistic signal is generated, either through photophysics, e.g., (direct) stochastic optical reconstruction microscopy ((d)STORM)^4,5^, photoactivated localization microscopy (PALM)^6^, or though binding, e.g., points accumulation for imaging in nanoscale topography (PAINT and DNA-PAINT)^7–9^. Regardless of the method, data from SMLM experiments may be represented as a list of Cartesian coordinates of all localised fluorophores. Thus, these data are suitable for the application of statistical methods that can identify characteristics about the spatial arrangement of such data^10^. One of the most common methods is cluster analysis^11^.

Clustering methods can be classified into two groups: global clustering and complete clustering. Global clustering analysis returns an ensemble result on whether a point pattern is clustered or not; Ripley’s K-function^12^, nearest neighbour analysis (NNA)^13^, and the pair correlation function (PCF)^14^ have all been applied to SMLM data^15–19^. For example, Ripley’s K-function has revealed that the T-cell adaptor protein, LAT is clustered at the immunological synapse^15,16^. Global approaches are statistically robust but provide a limited description of the data. Complete clustering approaches have therefore gained popularity. These assign every localisation to a specific cluster or into a non-clustered population^20–27^. Complete clustering methods provide rich descriptions of the data such as the number of clusters, cluster shapes and so on. For example, the density-based spatial clustering of applications with noise algorithm (DBSCAN)^28^ has been applied to observe the clustering of the T cell receptor at the immune synapse^29^, RNA polymerase organization in *E.coli*^30^, and dopamine receptor clusters in neurons^31^.

A consensus framework for assessing algorithms used to generate complete clusterings of SMLM data remains to be developed. Yet, there are existing mathematical means by which to assess the accuracy, or success, of clustering algorithms in general^32^ by comparing the result to a known ground truth, simulated data set. One such metric is the Rand Index (RI)^32^, which aims to determine which points have the same cluster classification in the ground truth and the analysed output. The Rand Index has been modified in order to account for correct classifications that may arise by chance^33^, resulting in the Adjusted Rand Index (ARI, **Figure 1a**). The ARI has a range between −1 and 1, with high positive values representing good clustering agreement (**Supplementary Figure 1a-c**). Performance can also be measured geometrically, for example, by taking the convex hull of points assigned to the cluster and measuring the overlap between the output and the ground truth. In this context, points at the edge of the identified cluster become more significant (**Figure 1b**). This is the basis for another accuracy metric, the intersection over union (IoU, **Figure 1b**)^34–36^ whose value runs from 0-1, 1 representing perfect overlap between cluster areas. As well as evaluating performance, these tests can serve as a basis for assessing the suitability of clustering algorithm parameters/settings and thus inform their most effective use.

**Figure 1.**
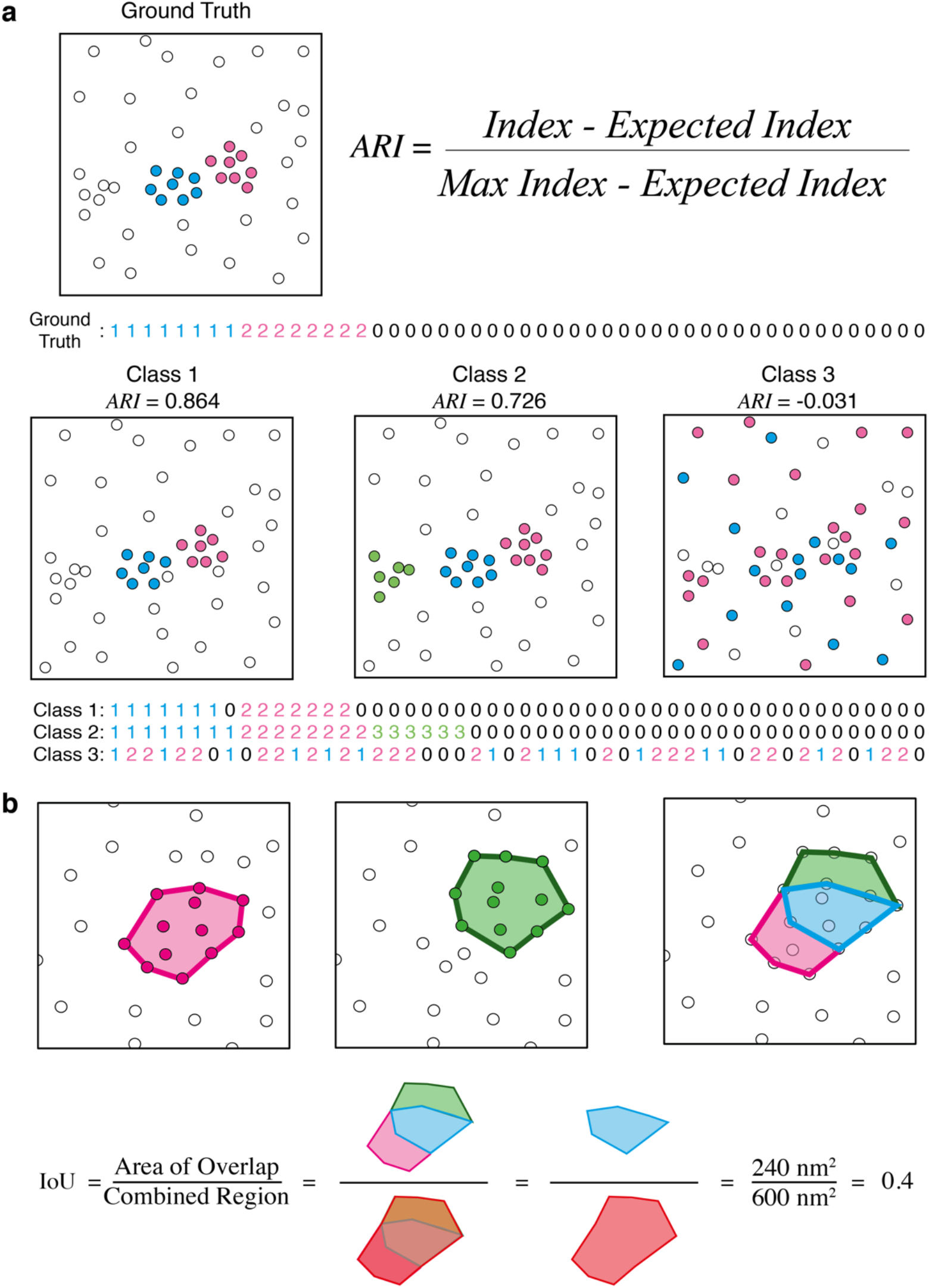
Clustering performance metrics. **a**) The Adjusted Rand Index (ARI) computes a measure of performance based on membership. **b**) The intersection over union (IoU) computes a measure of performance based on spatial overlap of clustered areas in the analysed data (green) and the ground truth (magenta).

Here, we implement ARI and IoU for the cluster analysis of simulated SMLM data mimicking common biological scenarios. To demonstrate the framework, we analyse the data with 3 different algorithms: DBSCAN, Topological Mode Analysis Tool (ToMATo), and kernel density estimation (KDE). We chose a range of different values for the user defined inputs, to uncover the optimal settings for these algorithms. Finally, we added a common experimental feature of SMLM: multiple blinking. We again assessed the algorithms in order to evaluate their robustness against this effect.

## Results

### Data simulation and pre-evaluation

Data generated from SMLM experiments can exhibit a wide range of spatial organisation, owing to the underlying biology and sample labelling. This may include varied levels of non-clustered points, sparse data, large numbers of clusters and irregular cluster shapes. These properties impact both the optimal algorithm parameters and their performance. We simulated 9 different ground truth molecule arrangements in 2000 × 2000 nm regions (**Table 1**, **Figure 2a-i, Online Methods**). *Scenario 1* - non-clustered single molecules seeded at completely spatially random positions (CSR; **Figure 2a**); *Scenario 2* - 20 clusters of 15 molecules per cluster with 50% of the total molecules being clustered (**Figure 2b**); *Scenario 3* - 20 clusters of 15 molecules per cluster with 20% of the total molecules being clustered (**Figure 2c**); *Scenario 4* - 20 clusters of 5 molecules per cluster with 50% of the total molecules being clustered (**Figure 2d**); *Scenario 5* - 100 clusters of 15 molecules per cluster with 50% of molecules being clustered (**Figure 2e**); *Scenario 6* - 20 elliptically shaped clusters aspect ratio 3:1 each of 50 molecules with 50% of the total molecules being clustered (**Figure 2f**); *Scenario 7* - 10 clusters with a distribution width of 25 nm and 10 clusters with a width of 75 nm, with 50% of the total molecules clustered (**Figure 2g**); *Scenario 8* - 10 clusters with 5 molecules per cluster and 10 molecules with 15 molecules per cluster, with 50% of the total molecules clustered (**Figure 2h**); *Scenario 9* - 10 clusters with 15 molecules per cluster, and a distribution with of 25 nm, and a further 10 clusters with 135 molecules and a distribution width of 75 nm, thus maintaining molecule density with increased size, with 50% of the total molecules clustered (**Figure 2i**). Clustering algorithms will produce erroneous clustering results when used on data that has little inherent clustering. Ripley’s K-function was computed for all of the scenarios above (**Figure 2a-i**, bottom panels). All but *Scenario 1* show significant deviation from a simulated curve for a completely random distribution (**Figure 2b-i**, bottom panels, red dotted line). Therefore, *Scenario 1* is not classified as significantly clustered using this approach, and hence not appropriate for further cluster identification.

**Table 1.** Properties of different simulated cluster scenarios. Parameters used for simulation of *Scenarios 1-9*. Red indicates deviations from *Scenario 2* for each scenario, whilst blue and red in *Scenario 9* indicate matching cluster parameters in that scenario.

| Simulation | No. clusters | Molecules per cluster | Elliptical | Cluster width ($\sigma$) | No. background molecules | % of single non-clustered molecules |
| --- | --- | --- | --- | --- | --- | --- |
| *Scenario 1* | 0 | N/A | N/A | 15 | 300 | 100 |
| *Scenario 2* | 20 | 15 | N | 15 | 300 | 50 |
| *Scenario 3* | 20 | 15 | N | 15 | 1500 | 80 |
| *Scenario 4* | 20 | 5 | N | 15 | 300 | 50 |
| *Scenario 5* | 100 | 15 | N | 15 | 1500 | 50 |
| *Scenario 6* | 20 | 50 | Y | x = 25,<br>y = 75 | 1000 | 50 |
| *Scenario 7* | 20 | 15 | N | 10 clusters = 25<br>10 clusters = 75 | 300 | 50 |
| *Scenario 8* | 20 | 10 clusters = 5<br>10 clusters = 15 | N | 15 | 200 | 50 |
| *Scenario 9* | 20 | 10 clusters = 15<br>10 clusters = 135 | N | 10 clusters = 25<br>10 clusters = 75 | 1500 | 50 |

**Figure 2 –.**
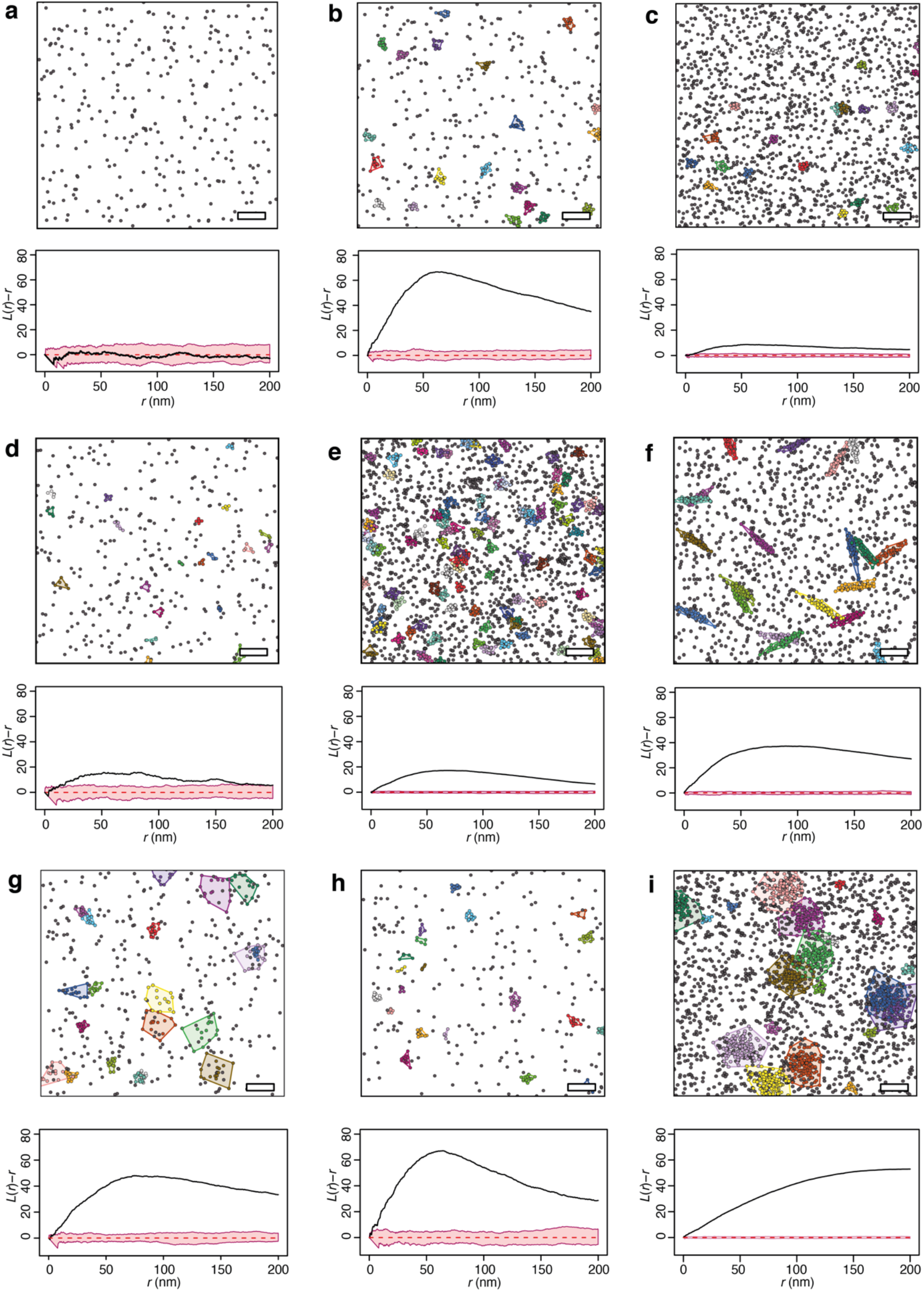
Examples of simulated data conditions and pre-evaluation of suitability. **a**) *Scenario 1* **b**) *Scenario 2*, **c**) *Scenario 3*, **d**) *Scenario 4*, **e**) *Scenario 5*, **f**) *Scenario 6*, **g**) *Scenario 7*, **h**) *Scenario 8*, and **i**) *Scenario 9*. Top panels - examples of simulation data, Bottom panels - linear representation of Ripley’s K function (*L*(*r*)-*r*), black line for simulation data, red line - mean from 100 simulations using the same number of molecules placed completely randomly, with 95% simulation envelopes (pink). Scale bars - 200 nm

### Performance metrics

We next analyzed the simulations with clustering algorithms and scored the results. Importantly, given the breadth of clustering algorithms and analyses available, selecting algorithms that differ in their approach, but require similar user guidance was important. To this end, we chose DBSCAN^28^ (**Supplementary Figure 2a**), ToMATo (**Supplementary Figure 2b**)^25^, and kernel density estimation (KDE, **Supplementary Figure 2c**). These algorithms were chosen on the basis that they differ in the method by which they identify cluster assignments but have comparable parameters to be set by the user (**Supplementary Figure 2**). For example, DBSCAN determines cluster and cluster edges by the number of neighbours for a given point above a threshold, whereas KDE uses an image-based approach where the pixel information represents the kernel density and points are clustered using a threshold. All three algorithms require two user-defined inputs to be set in order to function: minPts and ε for DBSCAN, a threshold on density mode persistence and search radius for ToMATo, and kernel size (σ) and density threshold for KDE. Clustering was performed for each algorithm by scanning the parameter combinations. This was repeated for 50 simulations within each scenario and the mean scores for each parameter combination calculated along with the variance. The optimal clustering parameters for each algorithm within each scenario and cluster metric are summarised in **Supplementary Table 1**.

For *Scenario 2* (**Fig. 3a**), the performance of the three algorithms is shown for every combination of user analysis settings (**Fig 3b-d**). The data shows a broad maximum of scores across the parameter space; however, a rapid drop-off and high-variance region exists for small-scale parameter settings, meaning if the optimal settings are not known, it is safer to err on the high-scale side. **Fig 3e** shows the maximum scores (*i.e*., for the best user settings) for each of the algorithm. The full performance analysis for all algorithms against all scenarios assessed by both metrics is shown in **Supplementary Figures 3-9**. The best score obtained from the optimal combination of input parameters in each case is shown in **Figure 4 and Supplementary Figure 10**. The best performing algorithm can depend on the scenario. With high levels of non-clustered points (*Scenario 3*, **Figure 4a**) or many clusters (*Scenario 5*, **Figure 4c** and **Supplementary Figure 5c**), for example, KDE maintains higher scores than the other algorithms (**Supplementary Figure 5c**). Furthermore, KDE seems to deal well with scenarios with variable cluster size and density, *i.e*., *Scenarios 7* and *9* (**Supplementary Figures 7 and 9**).

**Figure 3:**
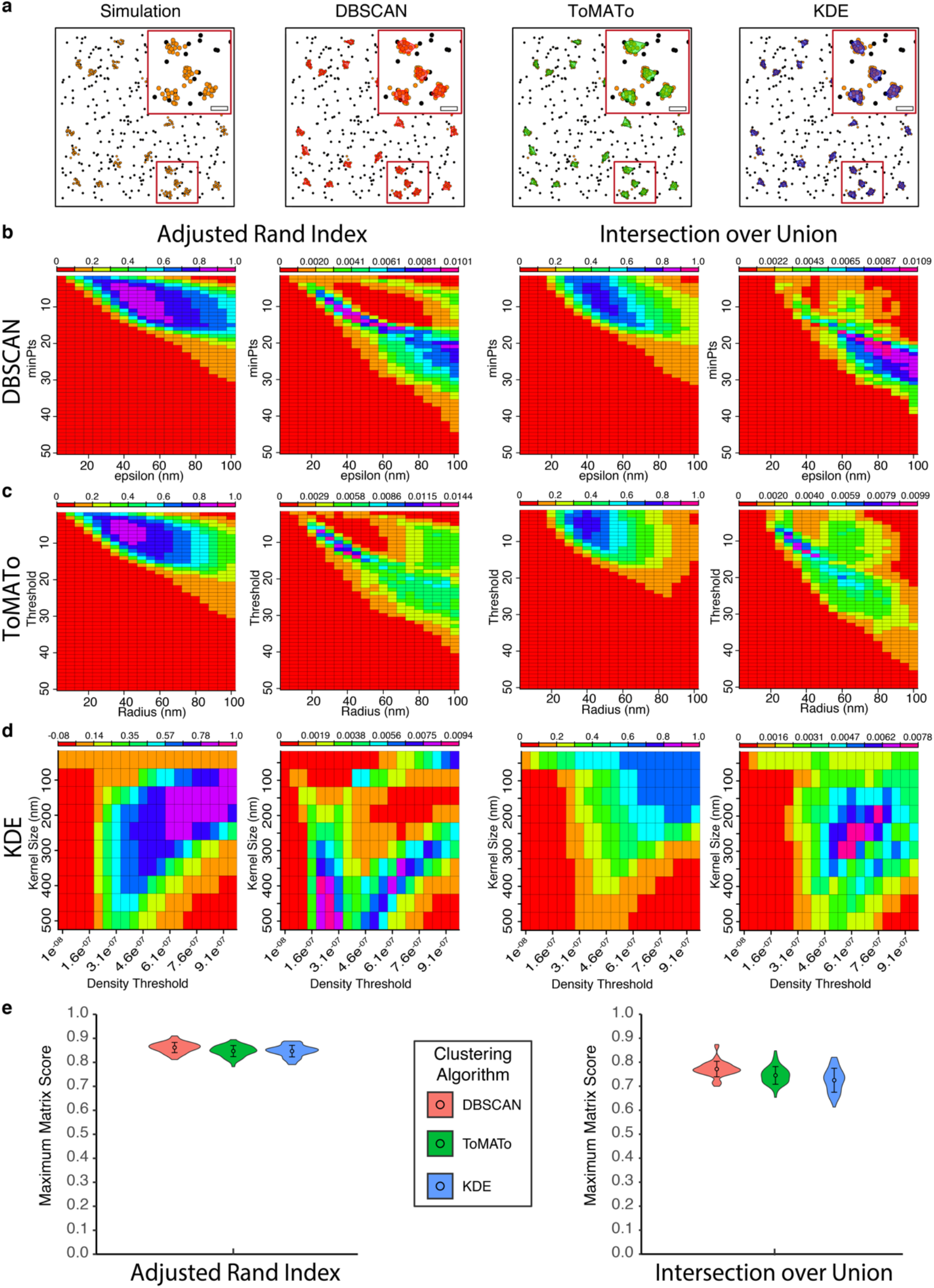
Performance analysis of the three algorithms against *Scenario 2*. **a**) Example of simulated data sets showing clustered points (orange) and cluster maps generated by DBSCAN (red), ToMATo (green), and KDE (blue). Zoom of single cluster for each algorithm (inset, scale bar - 100 nm) **b**) Mean and variance of the ARI and IoU scores for DBSCAN for all combinations of analysis parameters. **c**) Mean and variance of the ARI and IoU scores for ToMATo for all combinations of analysis parameters. **d**) Mean and variance of the ARI and IoU scores for KDE for all combinations of analysis parameters. **e**) Maximum ARI and IoU scores for the three algorithms with the mean and standard deviation.

**Figure 4.**
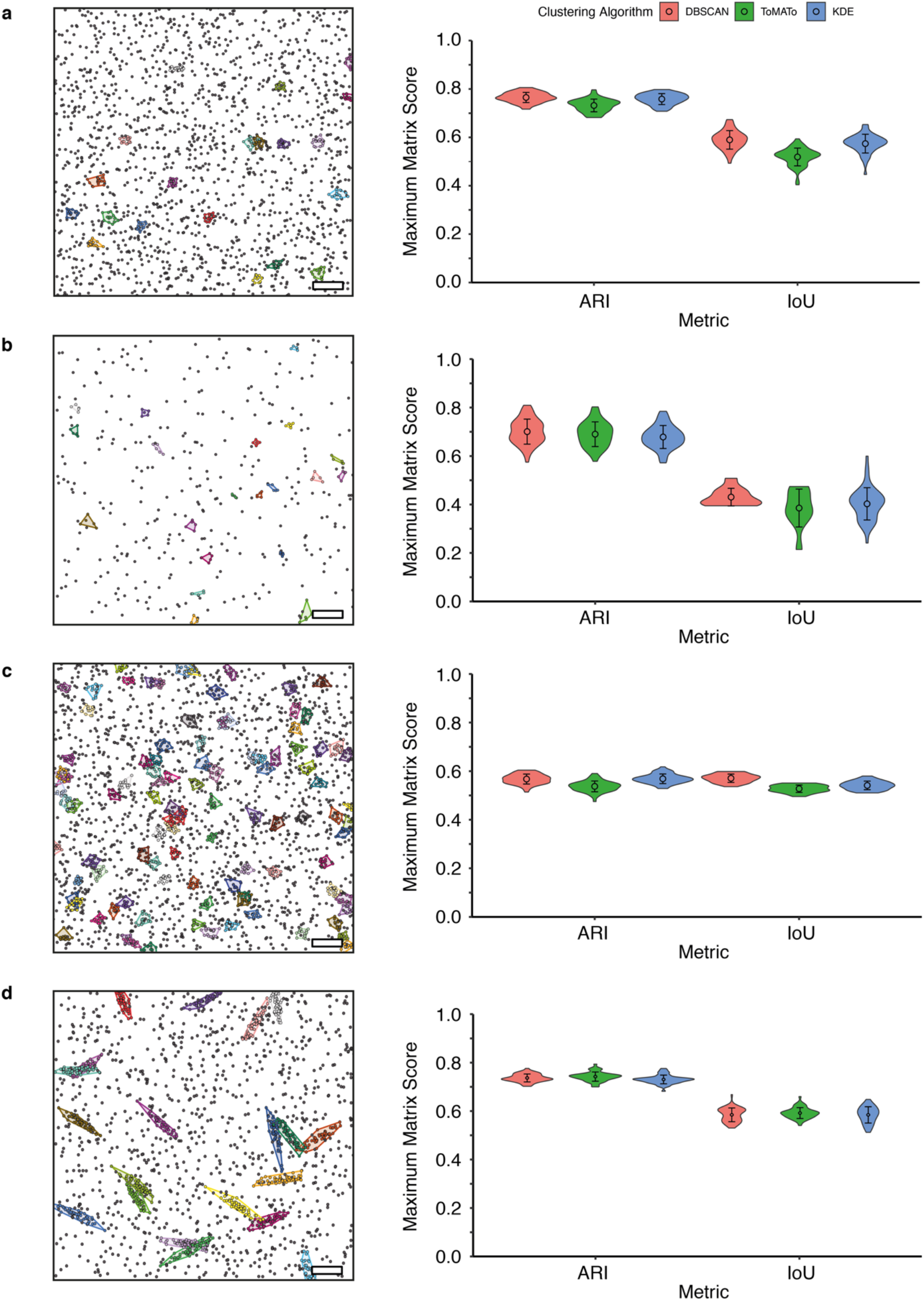
Performance of DBSCAN, ToMATo and KDE against Scenarios 3-6. Left: Representative ground truth clustering. Scale bar - 200 nm. Right: Maximum ARI and IoU scores; **a**) *Scenario 3*, **b**) *Scenario 4*, **c**) *Scenario 5*, **d**) *Scenario 6*, with the mean and standard deviation.

### Effect of multiple blinking on optimal clustering parameters

An inherent property of SMLM data is the presence of multiple points arising from a single molecule. In PALM data, approximate correction is possible by grouping localisations that appear close in space and time. These strategies are less for dSTORM data. Each ground truth fluorophore may therefore appear as a small cluster with membership related to the number of blinks and size related to the localisation precision. We tested the three cluster analysis algorithms’ performance against simulated data in the presence of multiple emission events. Each molecule in the ground truth conditions was assumed to be stoichiometrically labelled with a single fluorophore, and that the probability of that fluorophore blinking followed a geometric distribution^20^ set such that a single fluorophore would give 4-5 detections, on average. The optimal clustering parameters for each algorithm are summarised in **Supplementary Table 1**. The effect of this added blinking step on *Scenario 2* is shown in **Figure 5**, together with the complete performance evaluation for all three algorithms. There is an obvious drop off in performance for compared to the case with no blinking. The stochastic nature of the blinking introduces greater heterogeneity into the data. However, DBSCAN is more robust to the effect than the other two algorithms. KDE and ToMATo have similar performance by the ARI measure, but ToMATo is superior if the area of detected clusters is the required output (**Figure 5c**, right). This is a nice illustration that the choice of algorithm does not only depend on the nature of the underlying distribution, but also on the biological question being asked - *i.e*., which cluster descriptors are required to be accurate. The full performance analysis is shown in **Supplementary Figures 11-17** and the summary in **Figure 6** and **Supplementary Figure 18**.

**Figure 5:**
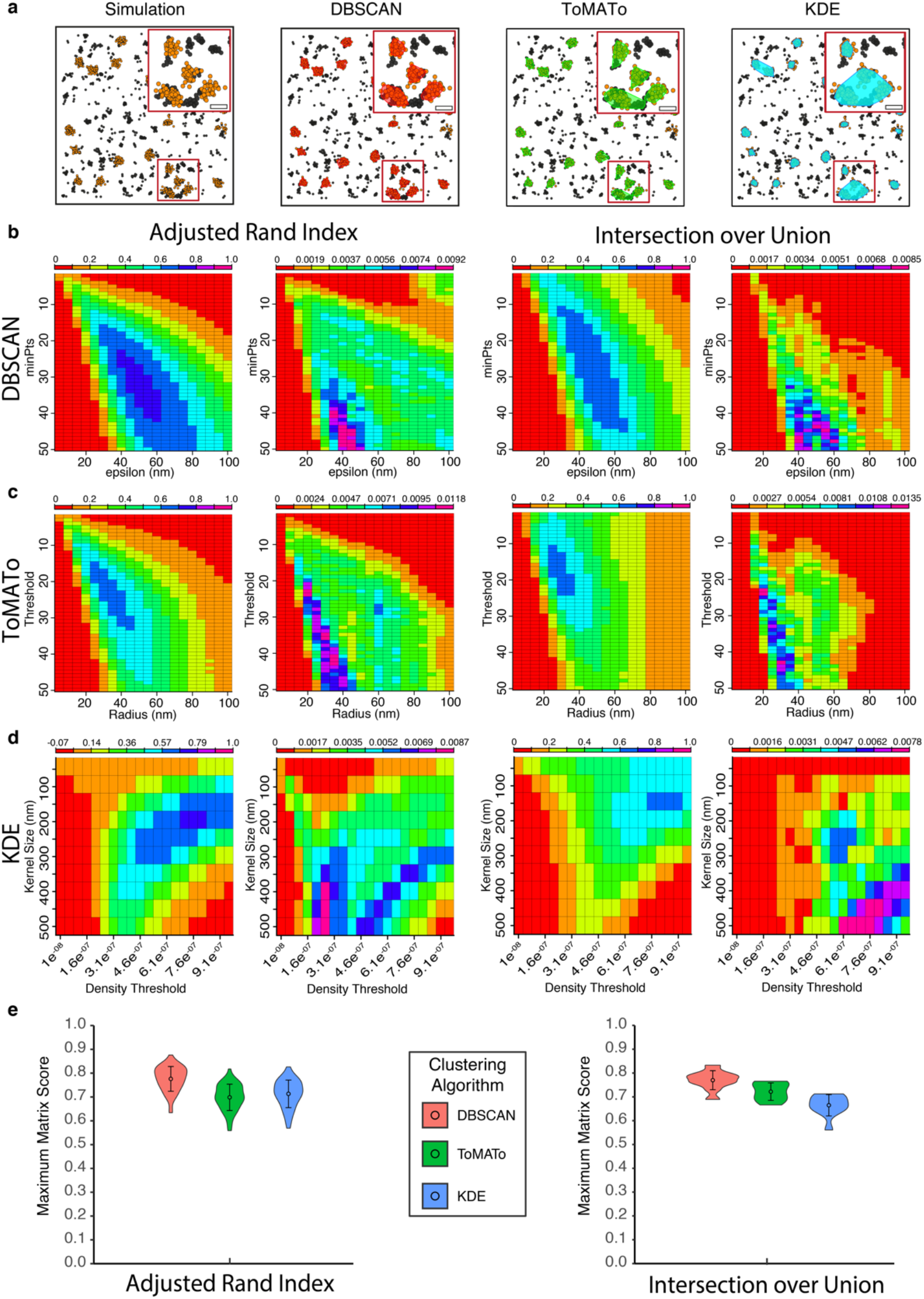
Performance against *Scenario 2* with the addition of fluorophore blinking. **a**) Example of simulated data sets showing the ground truth clusters (orange) and cluster maps generated by DBSCAN (red), ToMATo (green) and KDE (blue). **b**) Mean and variance of the ARI and IoU scores for DBSCAN for all combination is user analysis parameters. **c**) Mean and variance of the ARI and IoU scores for ToMATo for all combination is user analysis parameters. **d**) Mean and variance of the ARI and IoU scores for KDE for all combination is user analysis parameters. **e**) Maximum ARI and IoU scores for the three algorithms, with the mean and standard deviation.

**Figure 6.**
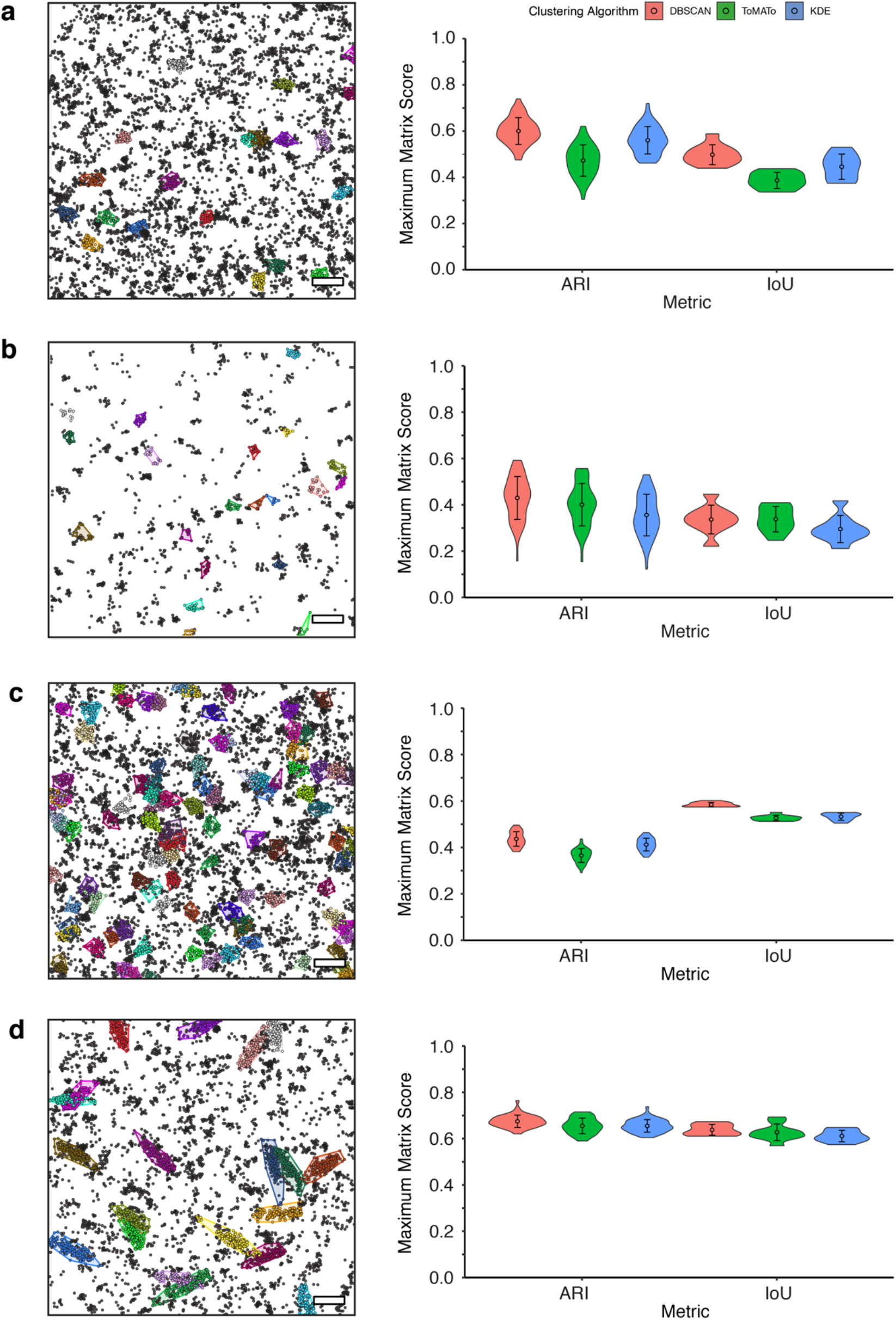
Performance of DBSCAN, ToMATo and KDE against *Scenarios 3-6* in the presence of multiple blinking. Left: Representative ground truth clustering showing localisations arising from clustered ground truth points. Right: Maximum ARI and IoU scores; **a**) *Scenario 3*, **b**) *Scenario 4*, **c**) *Scenario 5*, **d**) *Scenario 6*, with the mean and standard deviation.

## Discussion

SMLM produces data in the form of a pointillist set of localisation coordinates. A frequent goal is the cluster analysis of such data to extract quantitative information on molecular aggregation and to this end, a wide variety of algorithms have been deployed^11^. As development of new algorithms is ongoing, it is advantageous to have a framework in which the performance of these can be systematically evaluated. Here, we provide such an environment. Firstly, we propose a set of simulated point distributions with known ground truth clustering. We generated the same conditions with the presence of multiple blinking to provide more challenging conditions.

To assess the performance, we deployed two metrics designed for this purpose, ARI and IoU, which test different facets of the analysis performance. The ARI scores the analysis based on the similarity of cluster membership between the output and the ground truth and is computed based on whether every pair of points shared the same cluster label in each case. The input is therefore only the ordered set of cluster labels. IoU uses a measure of spatial overlap between the ground truth and detected cluster areas and takes as input the x,y coordinates as well as the labels. To illustrate the applicability of both metrics, we demonstrated them with the results from three cluster analysis algorithms - DBSCAN, ToMATo and KDE. These algorithms are diverse, but all require a choice of parameters by the user. We tested the performance of the algorithms over a range of parameter combinations and scored the results using ARI and IoU.

ARI typically returned higher scores than IoU for the majority of ground truth scenarios, without multiple blinking. This is because the convex hull for IoU relies on a small number of molecules at the periphery of the clusters. These are the points most likely to be misassigned by the algorithms. Thus, ARI is the most appropriate metric when clusters have a long tail to their molecule distribution. However, it is clear that the ARI metric operates best when the level of cluster overlap is low, and/or the size of the clusters is consistent as both these effects can lead to overlap of ground truth clusters. IoU is therefore a better measure of performance if heavy cluster overlap is expected. IoU is also more robust against multiple blinking than ARI due to the increased number of localizations along the edges of the clusters. Similarly, the emergence of new clusters, emanating from blinking monomers, adds new cluster indexes to the result classes thus decreasing the overall score, although the true clusters are classified correctly. Finally, since the IoU takes as input the actual x,y coordinates as well as the labels, the IoU metric itself is dependent on the cluster analysis and the underlying distribution. It may therefore be less useful to compare IoU scores between ROIs and scenarios than between algorithms.

We anticipate the framework presented here can be used to evaluate the performance of present cluster analysis algorithms designed for SMLM, and to inform the development of future methodologies. Further, the basis of both metrics, *i.e*., point classification and geometric overlap, are easily extendable to 3D, thus widening the applicability of this framework.

## Methods

### Ground Truth Cluster Simulations

Simulations of ground truth molecule point patterns were generated using scripts written in R. For simulations based on multivariate normal distributions, the cluster centres are randomly generated within a square field of interest (here, equivalent to 2×2 μm). At each centre, molecules are then placed around it according to the random multivariate distribution. For symmetrical clusters, a single value of 25 nm is used for the standard deviation of the multivariate distribution (unless stated otherwise, Table 1), whereas for the elliptically shaped clusters two different standard deviation values, for the minor and major axes, respectively, 25 and 75 nm, are used. Further for elliptical clusters, they a rotated by a random angle around the centre of mass of the generated cluster. Each cluster generated within the simulations possesses a unique “cluster index” value, *i.e*., molecules from the same cluster will have the same index value. Background molecules are given the index value of “0”. For each cluster scenario, 50 simulations were generated.

The different parameter values used to generate the simulations in this work are summarised in the table below;

### Fluorescence Blinking Cluster Simulations

The positions of molecules/fluorophores in the ground truth cluster scenarios were used as the basis for simulating data which has multiple blinking and detection precision inherent in SMLM. The simulateSTORM.r script from the RSMLM package (available at: https://github.com/JeremyPike/RSMLM) was used to generate the blinking simulations^25^. Briefly, transition between the fluorescent on- and off-state were modelled using a geometric distribution^20,25^ with probability of transition to the dark state set to 0.2, generating on average 4-5 fluorescent on-states, and thus, detections per molecule. Blinking was applied to all molecules in the simulations, thus single background molecules were also prone to blinking here. Detections owing to a single molecule will all be ascribed the index value of that molecule from the ground truth, e.g., if detections are generated from a ground truth molecule with a cluster index of “5”, then the detections too will retain the cluster index of “5”. The localisation uncertainty for each blinking event was determined using a normal distribution centered on the molecule position. Standard deviation for localisation uncertainty was set using a log-normal distribution with mean 2.8 and standard deviation 0.28^20^.

### DBSCAN clustering and parameter scanning

The density based spatial clustering of applications with noise (DBSCAN) algorithm was implemented in R using the dbscan R package. For DBSCAN there are two parameters; ε, which is the radius of search around each point, and minPts, which is the minimum number of neighbouring points within that radius for the point to be assigned to the cluster. Points within ε of clustered points but failing to fulfil minPts are designated the edge of the cluster. For DBSCAN parameter scanning, the ε (nm) and minPts threshold were varied. For epsilon values of from a minimum of 5 nm was used and stepped by 5 nm up to a maximum value of 100 nm (20 steps). For the minPts threshold a minimum value of 2 was used and stepped by 1 up to a maximum value of 50 (49 steps). Therefore, for each simulation, 980 total combinations of epsilon *vs* minPts threshold were performed, and the resulting indexing from each combination retained for further analysis.

### ToMATo clustering and parameter scanning

Topological Mode Analysis Tool (ToMATo) algorithm was implemented in R using the clusterTomato function from the RSMLM library (available at: https://github.com/JeremyPike/RSMLM)^25^. For ToMATo parameter scanning, the search radius (nm) and birth density threshold were varied. For the search radius, a minimum of 5 nm was used and stepped by 5 nm up to a maximum value of 100 nm (20 steps). For the birth density threshold, a minimum value of 2 was used and stepped by 1 up to a maximum value of 50 (49 steps). Therefore, for each simulation, 980 total combinations of search radius *vs* birth density threshold were performed, and the resulting indexing from each combination retained for further analysis.

### Kernel Density Estimation clustering and parameter scanning

Kernel density estimation (KDE) was performed using the kde2d function form the MASS R package. A 2D matrix was generated using the minimum and maximum dimensions from the simulation data, with each element in the matrix corresponding to a 1 nm^2^ region. The simulation coordinates are then convolved with a 2D Gaussian kernel within this 2D matrix, and densities within each of these 1 nm^2^ regions after convolution calculated. This 2D density matrix can then be thresholded according to a specific density value, and higher density regions above the cut off are considered “clustered”. These regions are then used to assign points from the real data into clusters. For KDE parameter scanning, the 2D Gaussian kernel width and the density threshold were varied. For the kernel width values of a minimum of 50 nm was used and stepped by 50 nm up to a maximum value of 500 nm (10 steps). For the density threshold a minimum value of 0.1e^−07^ was used and stepped by 0.5e^−07^ up to a maximum value of 0.1e^−06^ (10 steps). Therefore, for each simulation, 100 total combinations of kernel size vs density threshold were performed, and the resulting indexing from each combination retained for further analysis.

### Adjusted Rand Index Scoring

Cluster indexes results from the clustering and parameter scanning are used to compute the adjusted Rand Index (ARI). The ARI calculation was implemented in R using the function mclustcomp from the dbscan library (full ARIR script can be found here: https://github.com/DJ-Nieves/ARI-and-IoU-cluster-analysis-evaluation). Briefly, the Rand Index (RI) is a measure of the similarity between two sets of cluster indexes, and is calculated using;

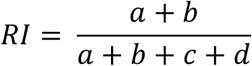

where *a* + *b* is the number of agreements between the ground truth and the cluster results, whereas *c* + *d* is number of disagreements between the ground truth and the cluster results.

The ARI corrects for the random chance of points being assigned to the correct clusters, and is calculated as follows;

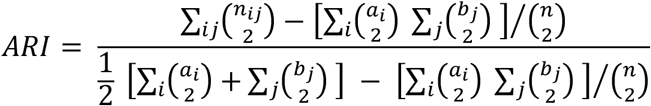

where *n_ij_* is the number of elements in common between clusterings *i* and *j*, *a_i_* is the sum of the contingency table for row *i*, and *b_j_* is the sum of the contingency table for column *j*.

For each parameter combination the clustering result (*i.e*., the cluster indexes) is compared to that of the ground truth simulation, and the ARI is calculated for that parameter combination. For fluorescent blinking simulations data, the clustering results are compared to the cluster indexes generated by the simulateSTORM algorithm as the “ground truth” condition. This is repeated for all parameter combinations to generate an ARI matrix.

### Intersection of Union Scoring

Intersection of union (IoU) scoring was implemented in custom written script in R (https://github.com/DJ-Nieves/ARI-and-IoU-cluster-analysis-evaluation). A convex hull was used to identify the molecule coordinates at the edge of each ground truth cluster. A filled polygon for each cluster was then generated within a binary image matching the limits of the data (pixel area = 1 nm^2^). All cluster images were then added together to generate a single image, and then the image was flattened to generate again a combined binary image. This process was performed for the ground truth clustering as well as each of the clustering results from the cluster parameter scanning. IoU is calculated as follows;

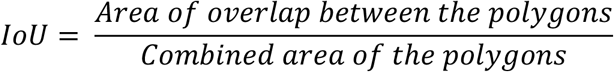

For the calculation from our binary images, the ground truth image was added to the cluster result image, thus giving a single image where the overlapping pixels had a value of 2. The number of pixels with value > 0 were equal to the combined area (nm^2^), whereas the number of pixels with value > 1 were equal to the area of overlap (nm^2^). For fluorescent blinking simulations data, the clustering results are compared to and image generated form the convex hulls of cluster indexes generated by the simulateSTORM algorithm as the “ground truth” condition. This is repeated for all parameter combinations to generate an IoU matrix.

## Data Availability

The simulation data used as the basis for this work is available for download at https://github.com/DJ-Nieves/ARI-and-IoU-cluster-analysis-evaluation without restriction. All other data are available upon request. R Code for calculating ARI and IoU for clustering results against a ground truth scenario is also available for download at https://github.com/DJ-Nieves/ARI-and-IoU-cluster-analysis-evaluation without restriction.

## Acknowledgements

D.O. acknowledges funding from BBSRC grant BB/R007365/1. MH acknowledges funding by the Deutsche Forschungsgemeinschaft (DFG, German Research Foundation, Project-ID 259130777, SFB 1177; GRK 2566).

## Author contributions

D.J.N., Wrote analysis code, performed analysis and wrote the manuscript. J.A.P. Wrote analysis code, F.L., J.G., D.S., E.A.K.C., A.A.P., J-B.S. and M.H. contributed ideas and concepts. D.M.O. Conceived the work and wrote the manuscript.

