## Supplementary for "A framework for evaluating the performance of SMLM cluster analysis algorithms"

### SUPPLEMENTARY FIGURES

**a**

Class 1

| Ground Truth | 0 | 1 | 2 |
|---|---|---|---|
| 0 | 34 | 0 | 0 |
| 1 | 1 | 7 | 0 |
| 2 | 1 | 0 | 7 |

$$\sum_{ij}\binom{n_{ij}}{2} = \binom{34}{2}+\binom{0}{2}+\binom{0}{2}+\binom{1}{2}+\binom{7}{2}+\binom{0}{2}+\binom{1}{2}+\binom{0}{2}+\binom{7}{2} = 561+0+0+0+21+0+0+0+21 = 603$$

$$\sum_{i}\binom{a_{i}}{2} = \binom{34}{2}+\binom{8}{2}+\binom{8}{2} = 561+28+28 = 617$$

$$\sum_{j}\binom{b_{j}}{2} = \binom{36}{2}+\binom{7}{2}+\binom{7}{2} = 630+21+21 = 672$$

$$ARI = \frac{603 - [617 \text{ x } 672]/\binom{50}{2}}{\frac{1}{2}[617+672] - [617 \text{ x } 672]/\binom{50}{2}} = 0.864393$$

**b**

Class 2

| Ground Truth | 0 | 1 | 2 | 3 |
|---|---|---|---|---|
| 0 | 28 | 0 | 0 | 6 |
| 1 | 0 | 8 | 0 | 0 |
| 2 | 0 | 0 | 8 | 0 |

$$\sum_{ij}\binom{n_{ij}}{2} = \binom{28}{2}+\binom{0}{2}+\binom{0}{2}+\binom{6}{2}+\binom{0}{2}+\binom{8}{2}+\binom{0}{2}+\binom{0}{2}+\binom{0}{2}+\binom{0}{2}+\binom{8}{2}+\binom{0}{2} = 378+0+0+15+0+28+0+0+0+0+28+0 = 449$$

$$\sum_{i}\binom{a_{i}}{2} = \binom{34}{2}+\binom{8}{2}+\binom{8}{2} = 561+28+28 = 617$$

$$\sum_{j}\binom{b_{j}}{2} = \binom{28}{2}+\binom{8}{2}+\binom{8}{2}+\binom{6}{2} = 378+28+28+15 = 449$$

$$ARI = \frac{449 - [617 \text{ x } 449]/\binom{50}{2}}{\frac{1}{2}[617+449] - [617 \text{ x } 449]/\binom{50}{2}} = 0.7262512$$

**c**

Class 3

| Ground Truth | 0 | 1 | 2 |
|---|---|---|---|
| 0 | 10 | 9 | 15 |
| 1 | 1 | 3 | 4 |
| 2 | 1 | 3 | 4 |

$$\sum_{ij}\binom{n_{ij}}{2} = \binom{10}{2}+\binom{9}{2}+\binom{15}{2}+\binom{1}{2}+\binom{3}{2}+\binom{4}{2}+\binom{1}{2}+\binom{3}{2}+\binom{4}{2} = 45+36+105+0+3+6+0+3+6 = 204$$

$$\sum_{i}\binom{a_{i}}{2} = \binom{34}{2}+\binom{8}{2}+\binom{8}{2} = 561+28+28 = 617$$

$$\sum_{j}\binom{b_{j}}{2} = \binom{12}{2}+\binom{15}{2}+\binom{23}{2} = 66+105+253 = 424$$

$$ARI = \frac{204 - [617 \text{ x } 424]/\binom{50}{2}}{\frac{1}{2}[617+424] - [617 \text{ x } 424]/\binom{50}{2}} = -0.03113793$$

**Supplementary Figure 1. ARI contingency tables and calculations for exemplar Classes in Figure 1a.** a) Ground truth vs Class 1, b) Ground truth vs Class 2, c) Ground truth vs Class 3.

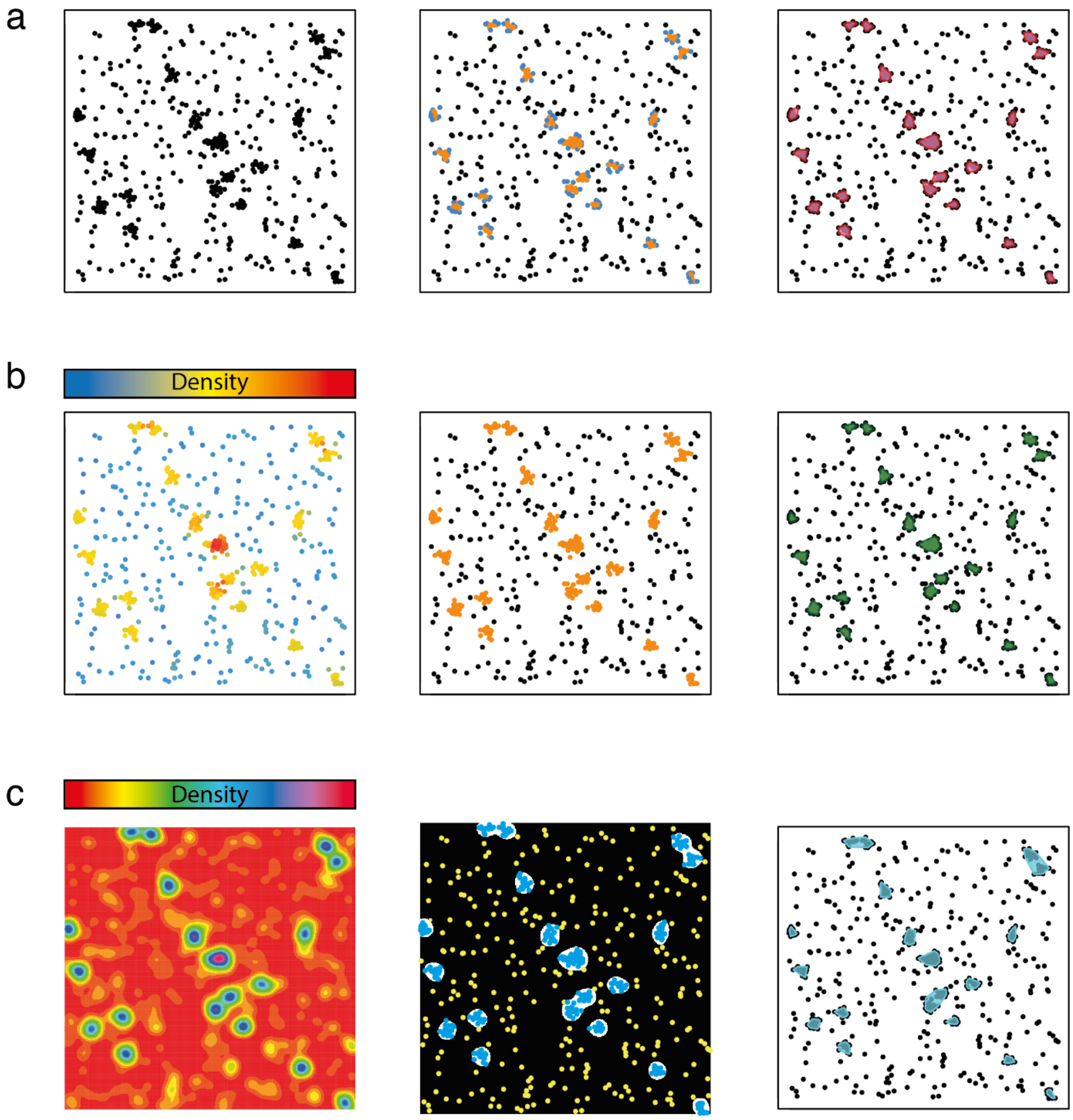


**Supplementary Figure 2. Clustering by DBSCAN, ToMATo and KDE.** a) The DBSCAN algorithm takes the coordinate data (left) and classifies the points according to two variables; ε, which is the radius of search around each point, and minPts, which is the minimum number of neighbouring points within that radius for the point of interest to be assigned to the cluster. In this example, ε = 60 nm, and minPts = 14. If a point satisfies the number of minPts within ε then it is classed as part of the cluster (middle, orange points) and points that do not satisfy minPts, but have cluster points within ε, are classed as edge points (middle, blue points). Coordinates that do not satisfy the minPts criteria and have no other coordinates within ε are classed as noise (middle, black points). This approach allows the segmentation of multiple clusters within the coordinate data signified by the magenta bounded polygons in the righthand result. b) The first step in ToMATo clustering is to calculate the density of points within the data and define local maxima, with high local density (left, yellow and red points). Points are assigned to local maxima (middle, orange) by following the gradient of the density. Candidate clusters are rejected, or merged, based on the strength, or persistence, of each local maximum.

In this implementation the extent to which this is done is dependent on a radius of search (35 nm) and threshold on density mode persistence (here, 5). Once complete, a clustering result is then produced (right green bounded polygons). c) KDE convolves each point within the coordinate data with a Gaussian function of defined width, here $\sigma = 150$ nm, and the summation of those Gaussians give the final image (left). The density, owing to the summation of multiple Gaussians, can be determined for each pixel in the image. This then allows thresholding of this image according to density/nm2 value (density threshold here was set to $7.6e^{-7}$). The thresholded image can be converted to a binary image, whereby the pixels above the threshold (middle, white) can now act as regions for segmenting the coordinate data. Points that fall within the white regions are defined as clustered (middle, blue) and those outside are designated as noise points (middle yellow). This produces a final cluster result (right, blue bounded polygons)

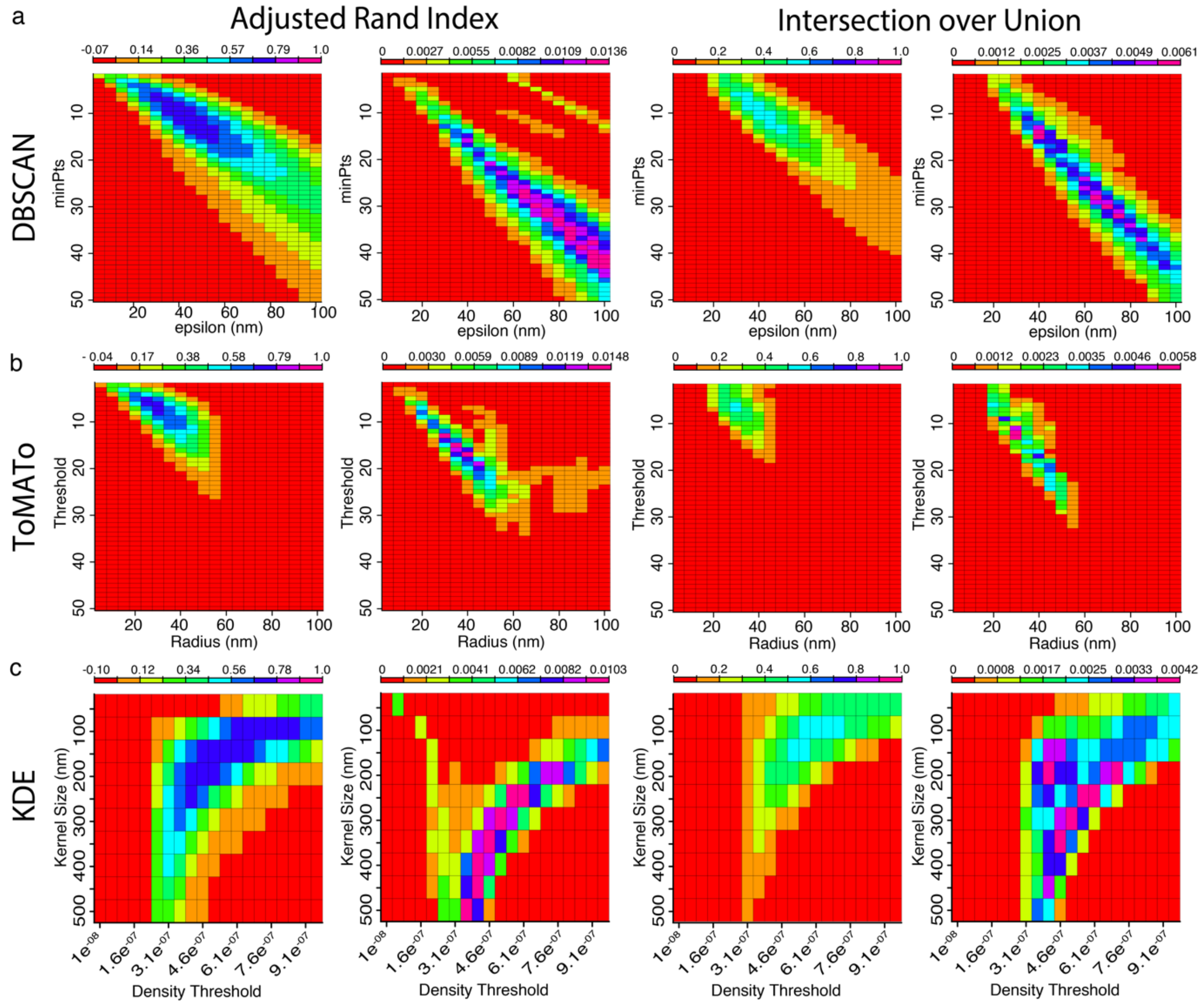


**Supplementary Figure 3. Parameter scan ARI and IoU matrices for DBSCAN, ToMATo, and KDE of Gaussian clusters with 80% background molecules added (*Scenario 3*).** a) Parameter scan matrices for DBSCAN assessed by the ARI (left) and IoU (right). For each index we have the mean index value matrix, i.e., the mean index value from the 50 simulations for each parameter combination (left), and the variance matrix, which is the variance of the mean index matrix values (right). b) Parameter scan matrices for ToMATo assessed by the ARI (left) and IoU (right), with the mean index value matrix (left), and the variance matrix (right). c) Parameter scan matrices for KDE assessed by the ARI (left) and IoU (right), with the mean index value matrix (left), and the variance matrix (right).

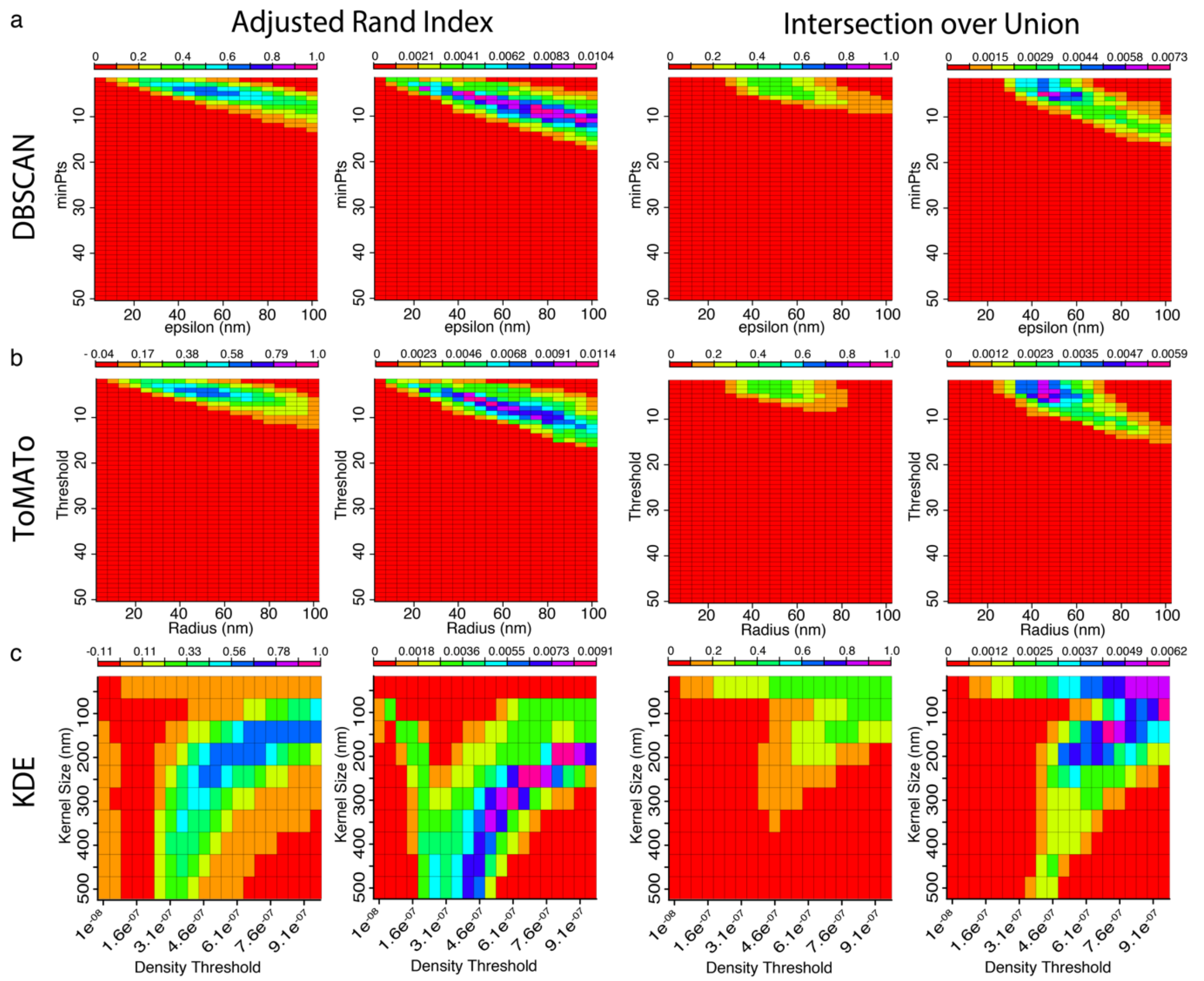


**Supplementary Figure 4. Parameter scan ARI and IoU matrices for DBSCAN, ToMATo, and KDE of low-density of molecules in Gaussian clusters (*Scenario 4*).** a) Parameter scan matrices for DBSCAN assessed by the ARI (left) and IoU (right). For each index we have the mean index value matrix, i.e., the mean index value from the 50 simulations for each parameter combination (left), and the variance matrix, which is the variance of the mean index matrix values (right). b) Parameter scan matrices for ToMATo assessed by the ARI (left) and IoU (right), with the mean index value matrix (left), and the variance matrix (right). c) Parameter scan matrices for KDE assessed by the ARI (left) and IoU (right), with the mean index value matrix (left), and the variance matrix (right).

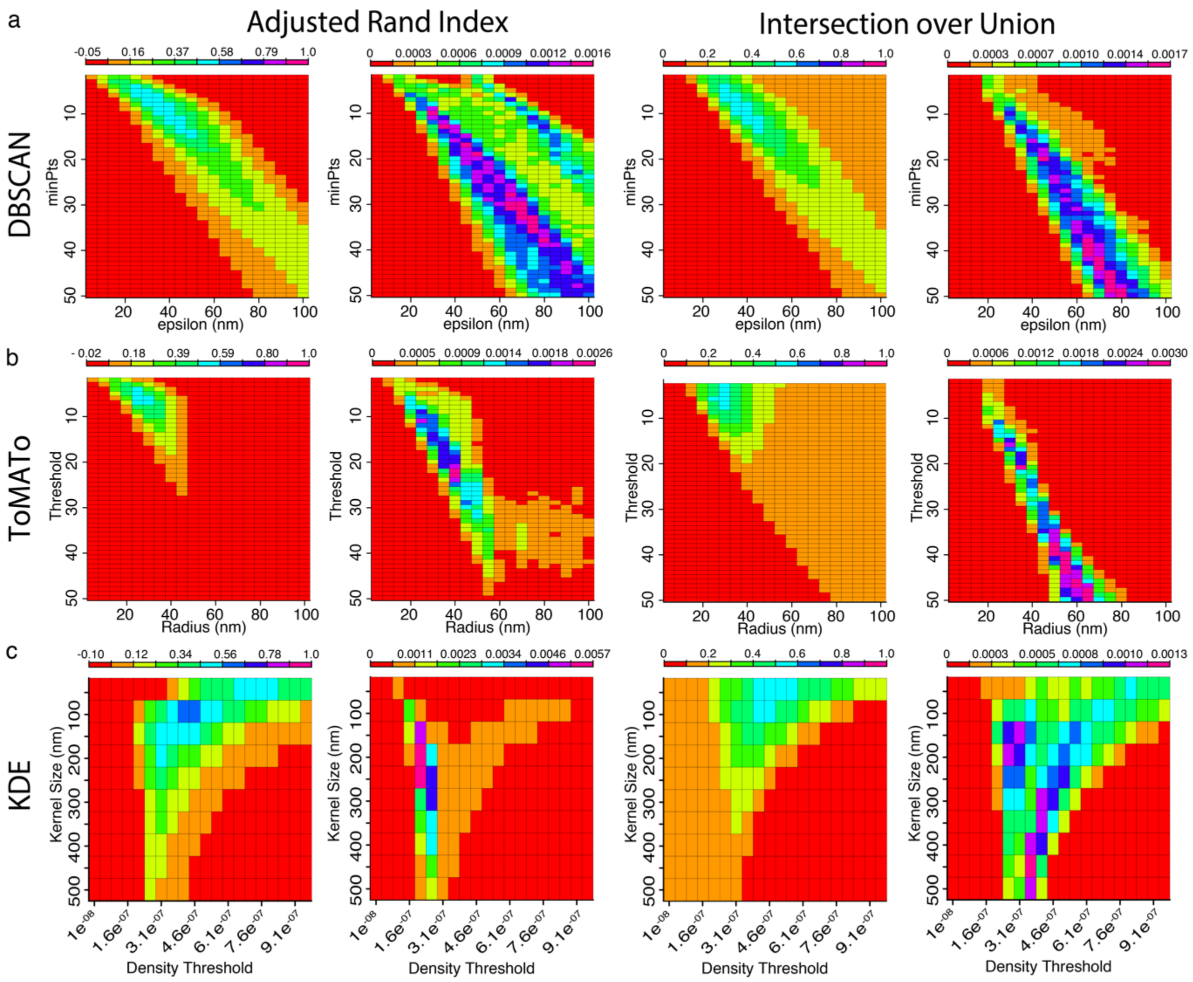


**Supplementary Figure 5. Parameter scan ARI and IoU matrices for DBSCAN, ToMATo, and KDE of high density of Gaussian clusters within the simulation area (*Scenario 5*).** a) Parameter scan matrices for DBSCAN assessed by the ARI (left) and IoU (right). For each index we have the mean index value matrix, i.e., the mean index value from the 50 simulations for each parameter combination (left), and the variance matrix, which is the variance of the mean index matrix values (right). b) Parameter scan matrices for ToMATo assessed by the ARI (left) and IoU (right), with the mean index value matrix (left), and the variance matrix (right). c) Parameter scan matrices for KDE assessed by the ARI (left) and IoU (right), with the mean index value matrix (left), and the variance matrix (right).

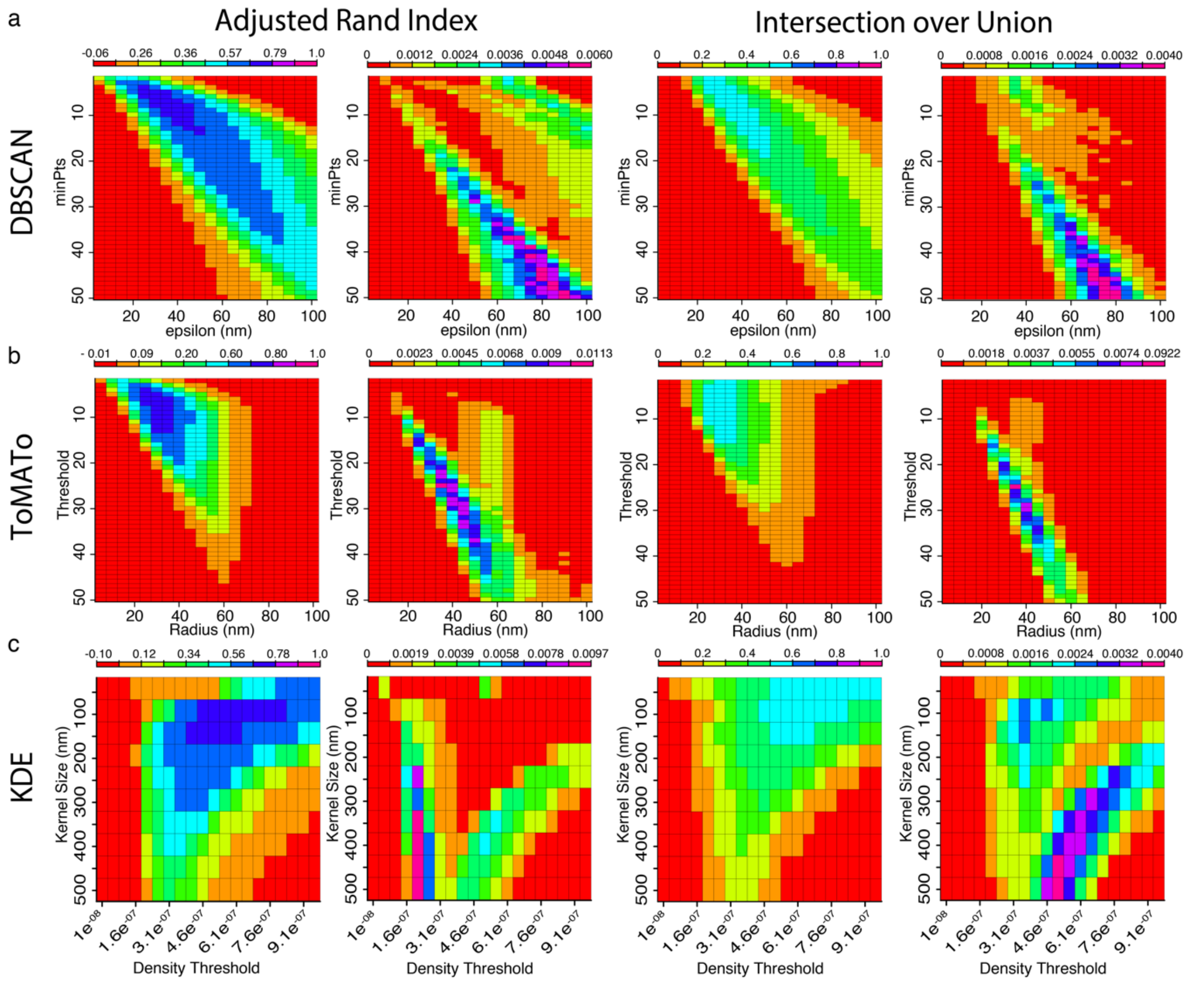


**Supplementary Figure 6. Parameter scan ARI and IoU matrices for DBSCAN, ToMATo, and KDE of elliptically shaped clusters (*Scenario 6*).** a) Parameter scan matrices for DBSCAN assessed by the ARI (left) and IoU (right). For each index we have the mean index value matrix, i.e., the mean index value from the 50 simulations for each parameter combination (left), and the variance matrix, which is the variance of the mean index matrix values (right). b) Parameter scan matrices for ToMATo assessed by the ARI (left) and IoU (right), with the mean index value matrix (left), and the variance matrix (right). c) Parameter scan matrices for KDE assessed by the ARI (left) and IoU (right), with the mean index value matrix (left), and the variance matrix (right).

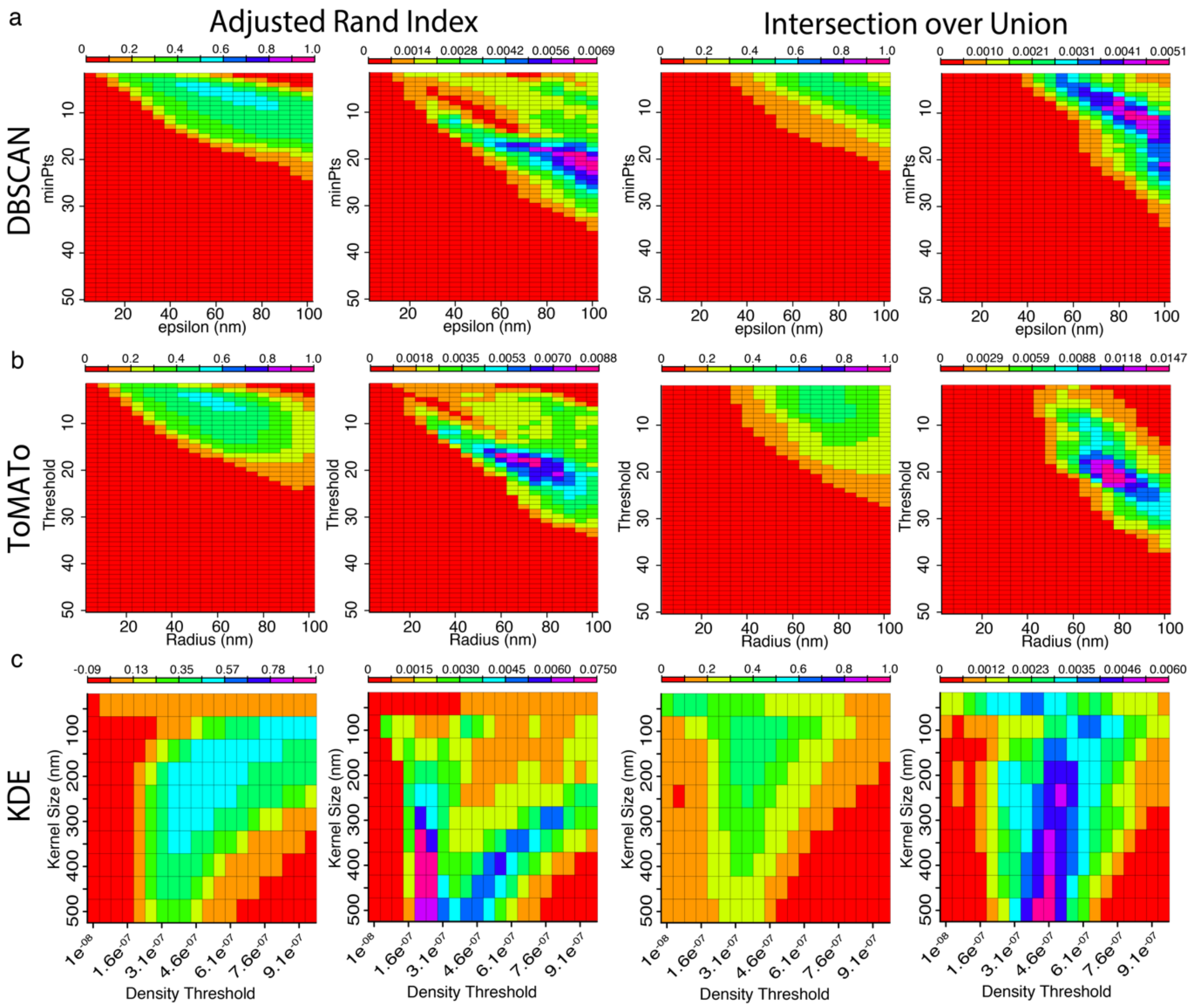


**Supplementary Figure 7. Parameter scan ARI and IoU matrices for DBSCAN, ToMATo, and KDE of clusters with different distribution widths (*Scenario 7*).** a) Parameter scan matrices for DBSCAN assessed by the ARI (left) and IoU (right). For each index we have the mean index value matrix, i.e., the mean index value from the 50 simulations for each parameter combination (left), and the variance matrix, which is the variance of the mean index matrix values (right). b) Parameter scan matrices for ToMATo assessed by the ARI (left) and IoU (right), with the mean index value matrix (left), and the variance matrix (right). c) Parameter scan matrices for KDE assessed by the ARI (left) and IoU (right), with the mean index value matrix (left), and the variance matrix (right).

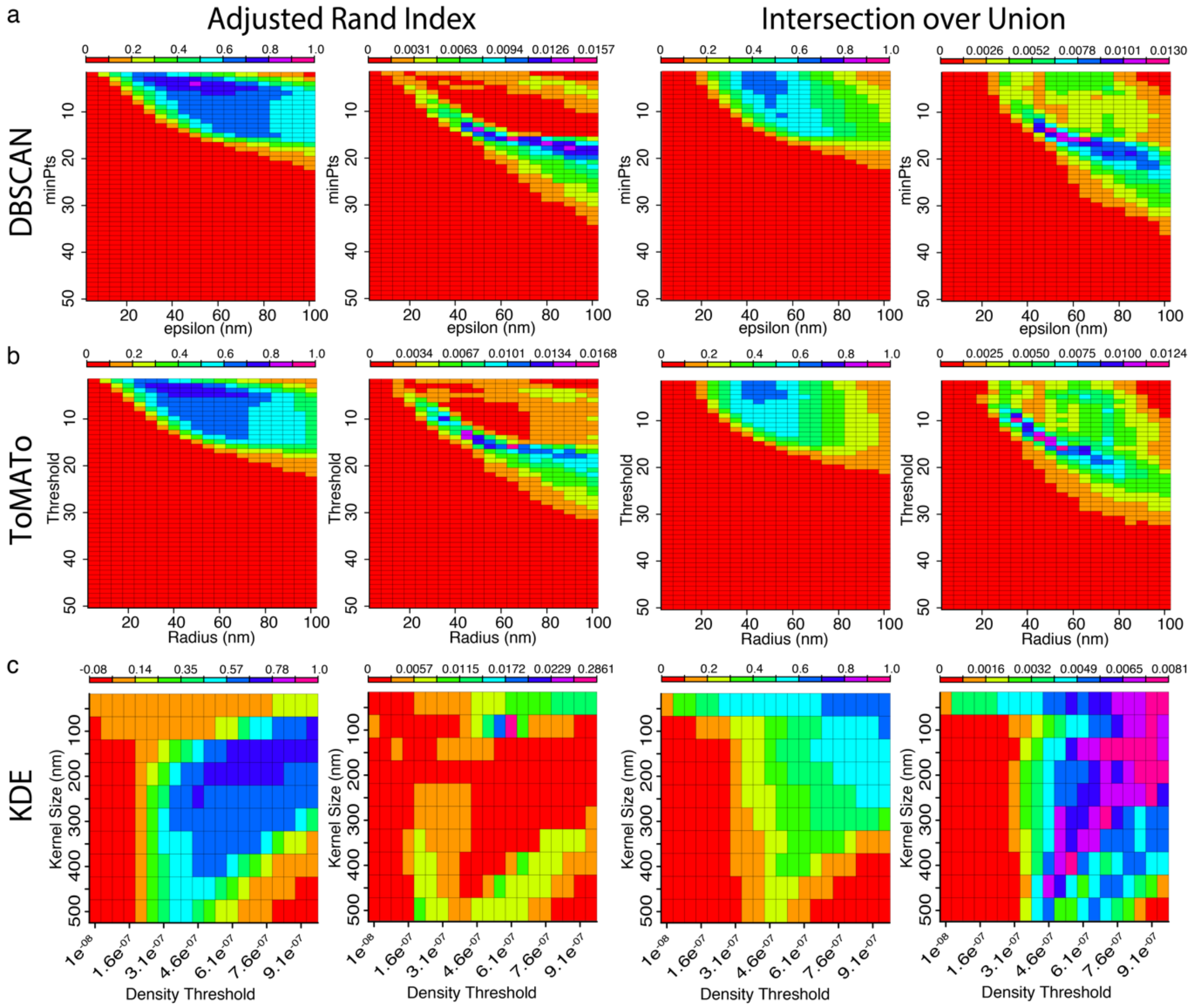


**Supplementary Figure 8. Parameter scan ARI and IoU matrices for DBSCAN, ToMATo, and KDE of clusters with different densities (*Scenario 8*).** a) Parameter scan matrices for DBSCAN assessed by the ARI (left) and IoU (right). For each index we have the mean index value matrix, i.e., the mean index value from the 50 simulations for each parameter combination (left), and the variance matrix, which is the variance of the mean index matrix values (right). b) Parameter scan matrices for ToMATo assessed by the ARI (left) and IoU (right), with the mean index value matrix (left), and the variance matrix (right). c) Parameter scan matrices for KDE assessed by the ARI (left) and IoU (right), with the mean index value matrix (left), and the variance matrix (right).

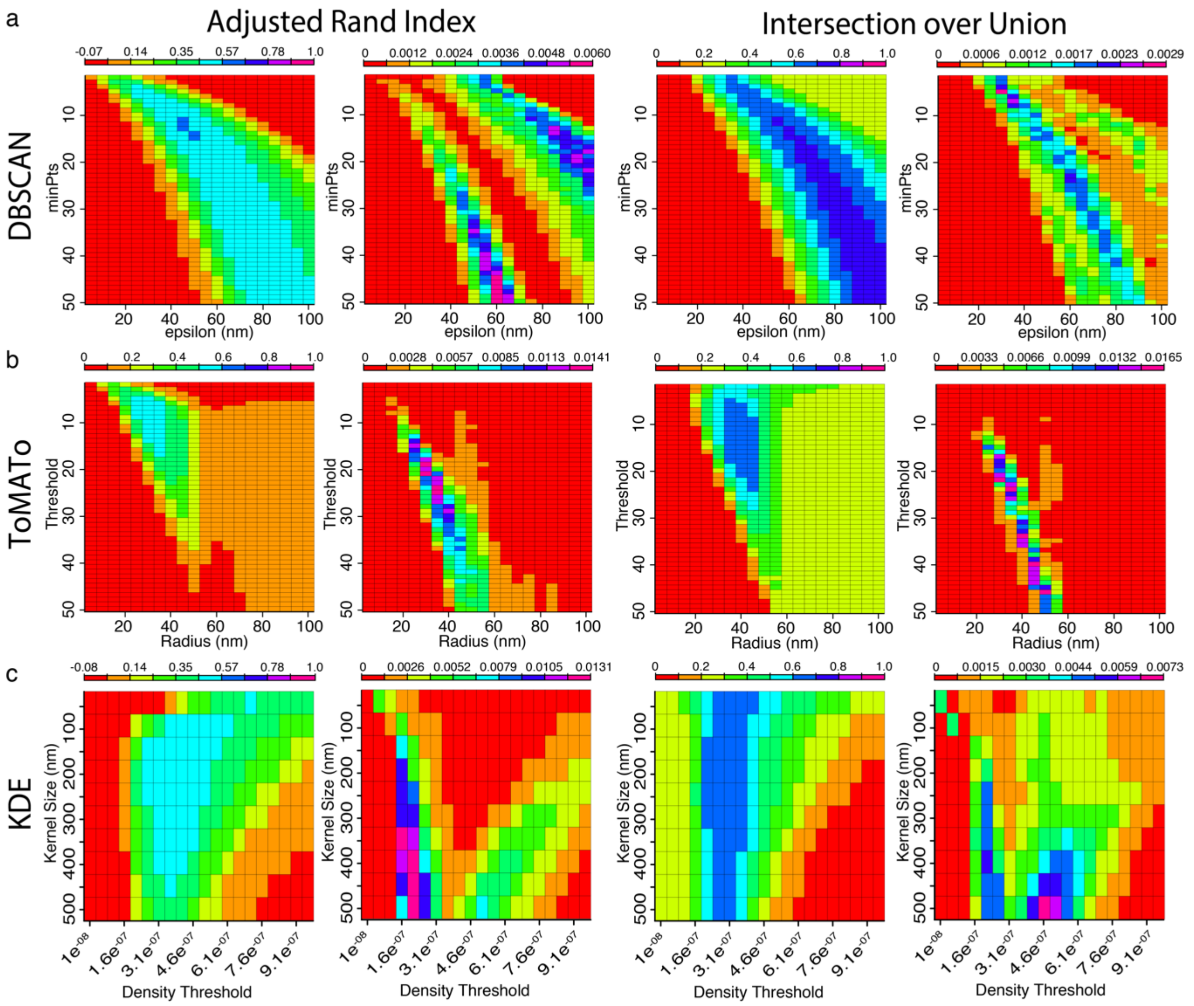


**Supplementary Figure 9. Parameter scan ARI and IoU matrices for DBSCAN, ToMATo, and KDE of clusters with different widths, but density maintained (*Scenario 9*).** a) Parameter scan matrices for DBSCAN assessed by the ARI (left) and IoU (right). For each index we have the mean index value matrix, i.e., the mean index value from the 50 simulations for each parameter combination (left), and the variance matrix, which is the variance of the mean index matrix values (right). b) Parameter scan matrices for ToMATo assessed by the ARI (left) and IoU (right), with the mean index value matrix (left), and the variance matrix (right). c) Parameter scan matrices for KDE assessed by the ARI (left) and IoU (right), with the mean index value matrix (left), and the variance matrix (right).

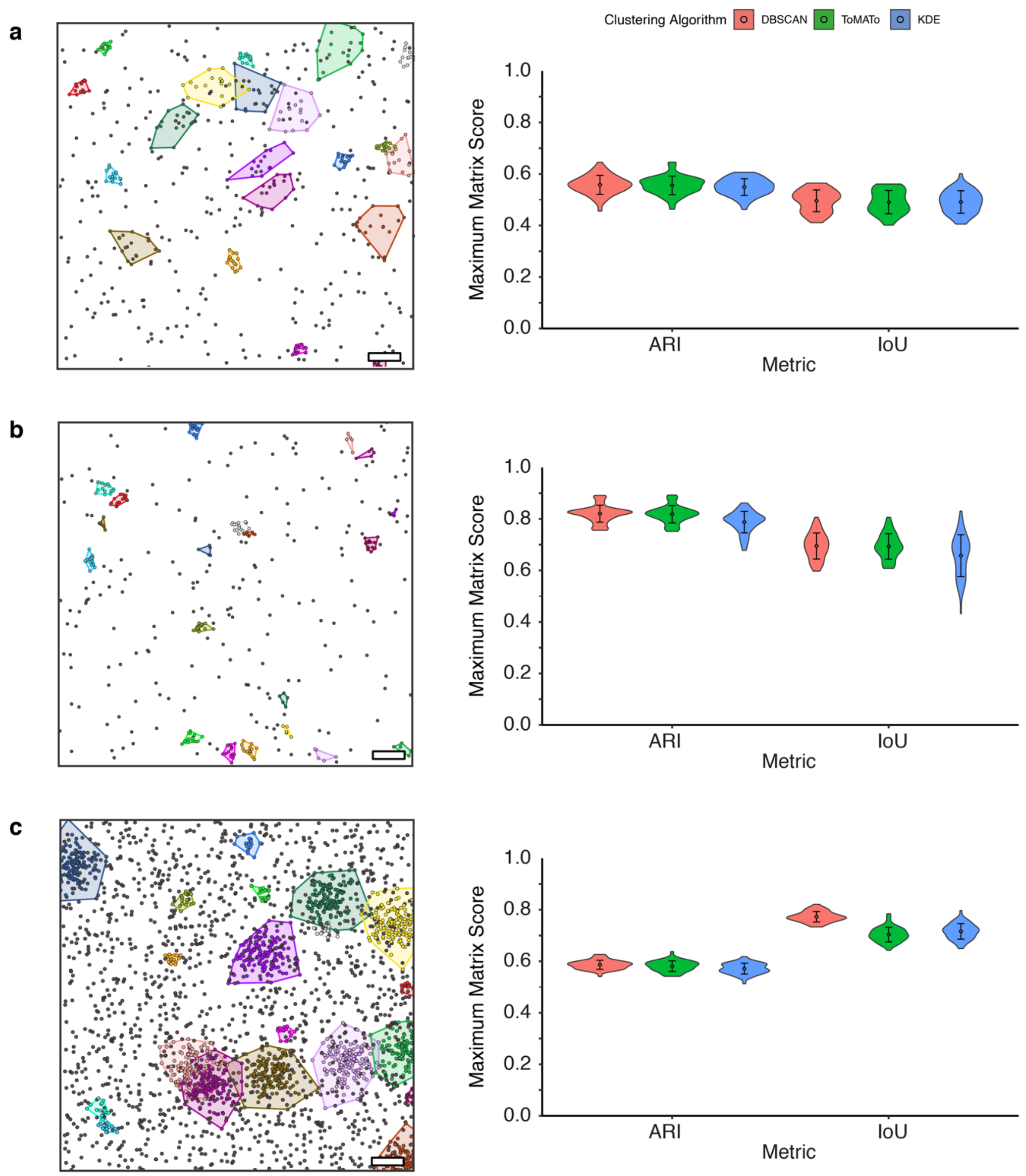


**Supplementary Figure 10. Performance of DBSCAN, ToMATo and KDE against Scenarios 7-9.** Left: Representative ground truth clustering showing clustered points (multicolor with bounding polygons) and non-clustered localisations (black). Scale bar – 200 nm. Right: Maximum ARI and IoU scores; **a)** *Scenario 7*, **b)** *Scenario 8*, **c)** *Scenario 9*,

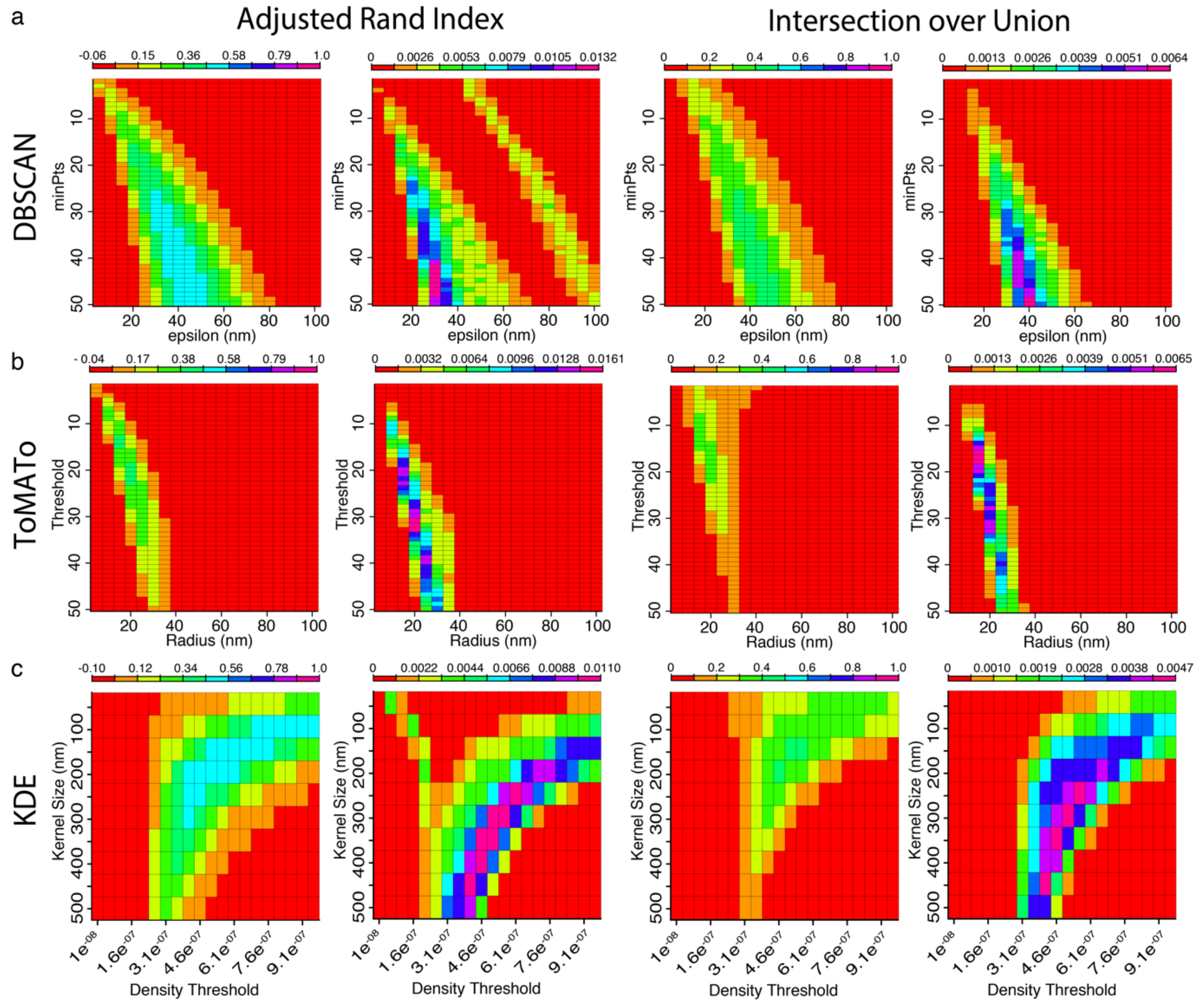


**Supplementary Figure 11. Parameter scan ARI and IoU matrices for DBSCAN, ToMATo, and KDE of Gaussian clusters with 80% background molecules, where fluorophore multiple blinking has been added (*Scenario 3*).** a) Parameter scan matrices for DBSCAN assessed by the ARI (left) and IoU (right). For each index we have the mean index value matrix, i.e., the mean index value from the 50 simulations for each parameter combination (left), and the variance matrix, which is the variance of the mean index matrix values (right). b) Parameter scan matrices for ToMATo assessed by the ARI (left) and IoU (right), with the mean index value matrix (left), and the variance matrix (right). c) Parameter scan matrices for KDE assessed by the ARI (left) and IoU (right), with the mean index value matrix (left), and the variance matrix (right).

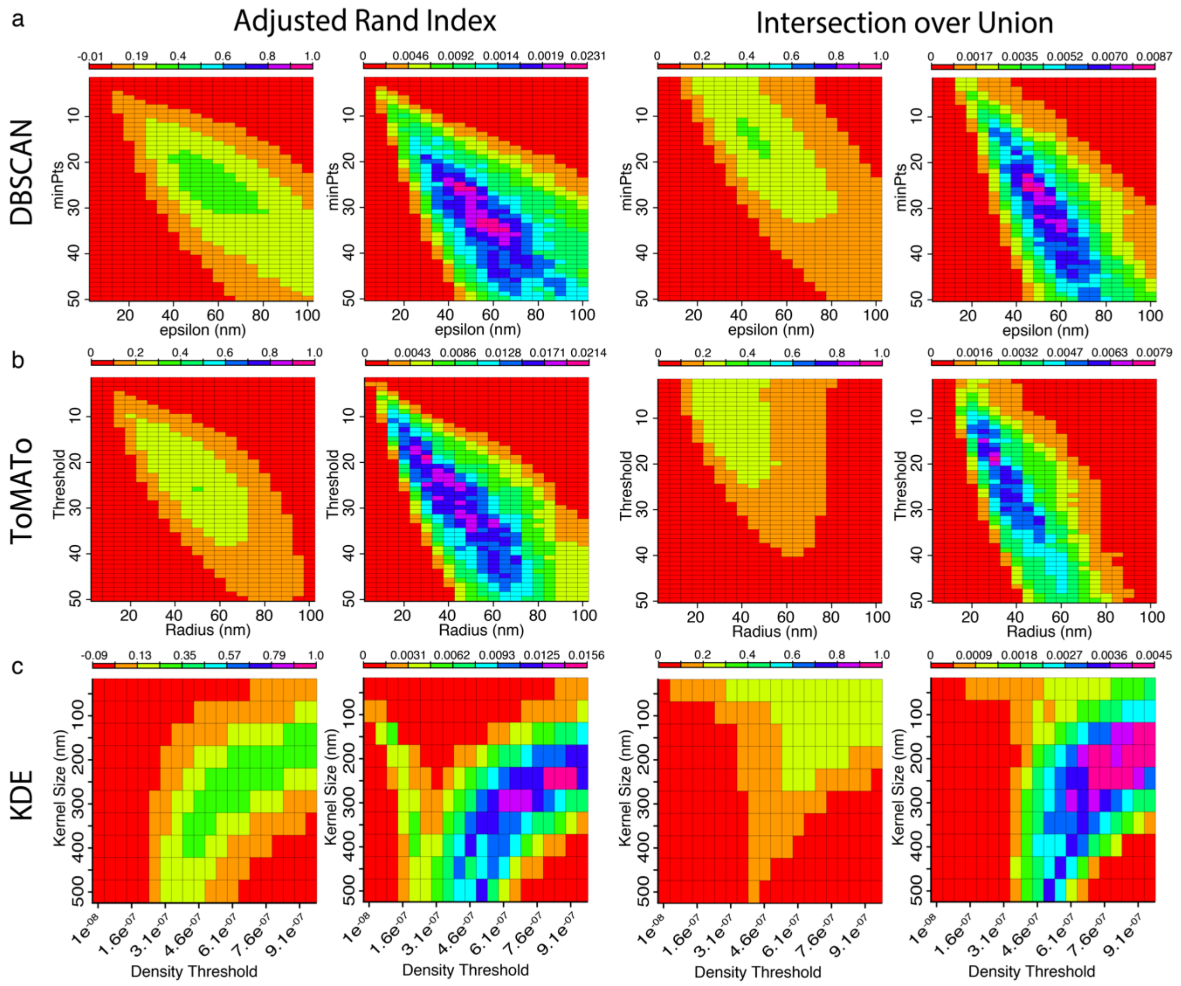


**Supplementary Figure 12. Parameter scan ARI and IoU matrices for DBSCAN, ToMATo, and KDE of low-density of molecules in Gaussian clusters with fluorophore multiple blinking added (*Scenario 4*).** a) Parameter scan matrices for DBSCAN assessed by the ARI (left) and IoU (right). For each index we have the mean index value matrix, i.e., the mean index value from the 50 simulations for each parameter combination (left), and the variance matrix, which is the variance of the mean index matrix values (right). b) Parameter scan matrices for ToMATo assessed by the ARI (left) and IoU (right), with the mean index value matrix (left), and the variance matrix (right). c) Parameter scan matrices for KDE assessed by the ARI (left) and IoU (right), with the mean index value matrix (left), and the variance matrix (right).

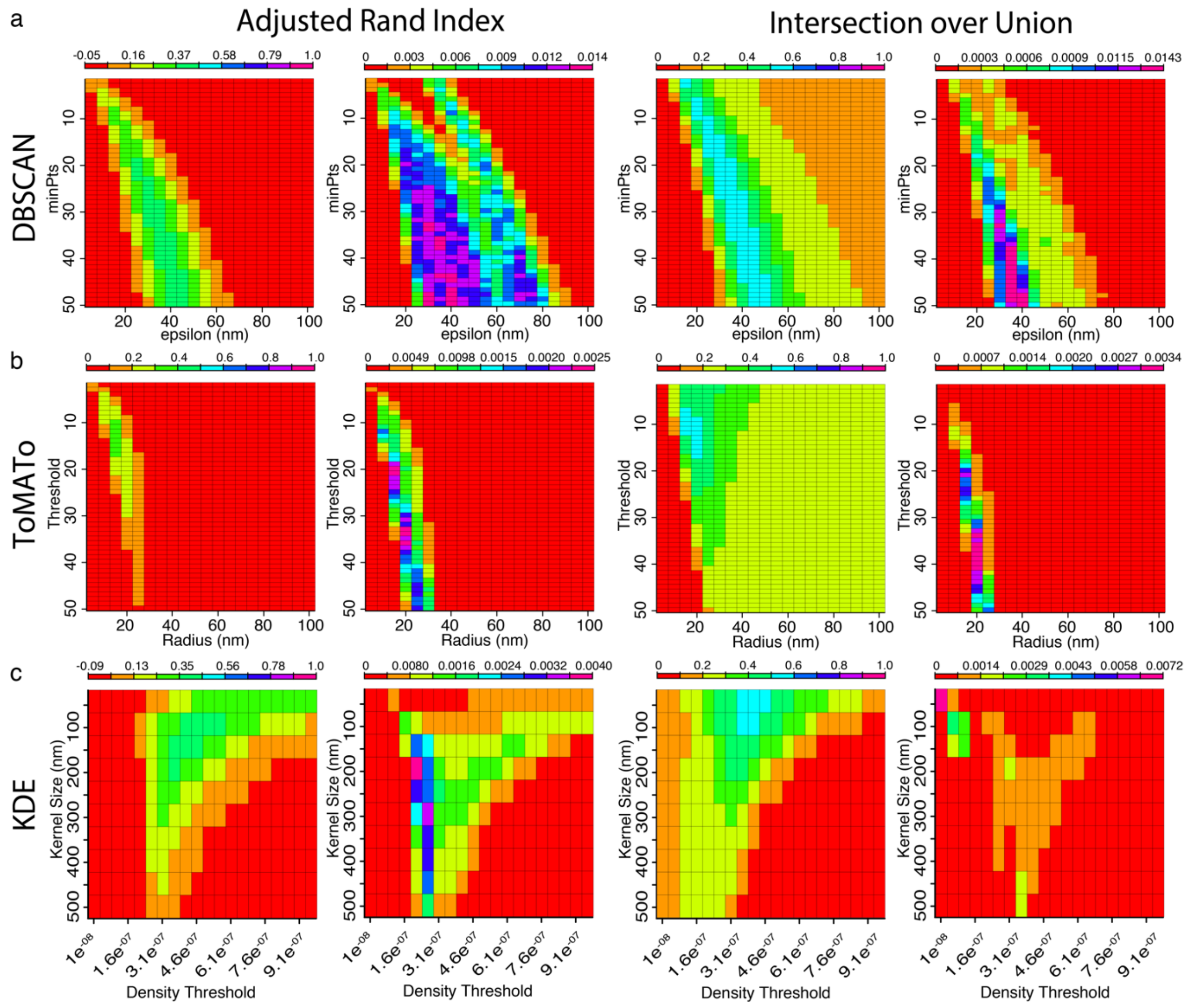


**Supplementary Figure 13. Parameter scan ARI and IoU matrices for DBSCAN, ToMATo, and KDE of high density of Gaussian clusters within the simulation area, with added fluorophore multiple blinking (*Scenario 5*).** a) Parameter scan matrices for DBSCAN assessed by the ARI (left) and IoU (right). For each index we have the mean index value matrix, i.e., the mean index value from the 50 simulations for each parameter combination (left), and the variance matrix, which is the variance of the mean index matrix values (right). b) Parameter scan matrices for ToMATo assessed by the ARI (left) and IoU (right), with the mean index value matrix (left), and the variance matrix (right). c) Parameter scan matrices for KDE assessed by the ARI (left) and IoU (right), with the mean index value matrix (left), and the variance matrix (right).

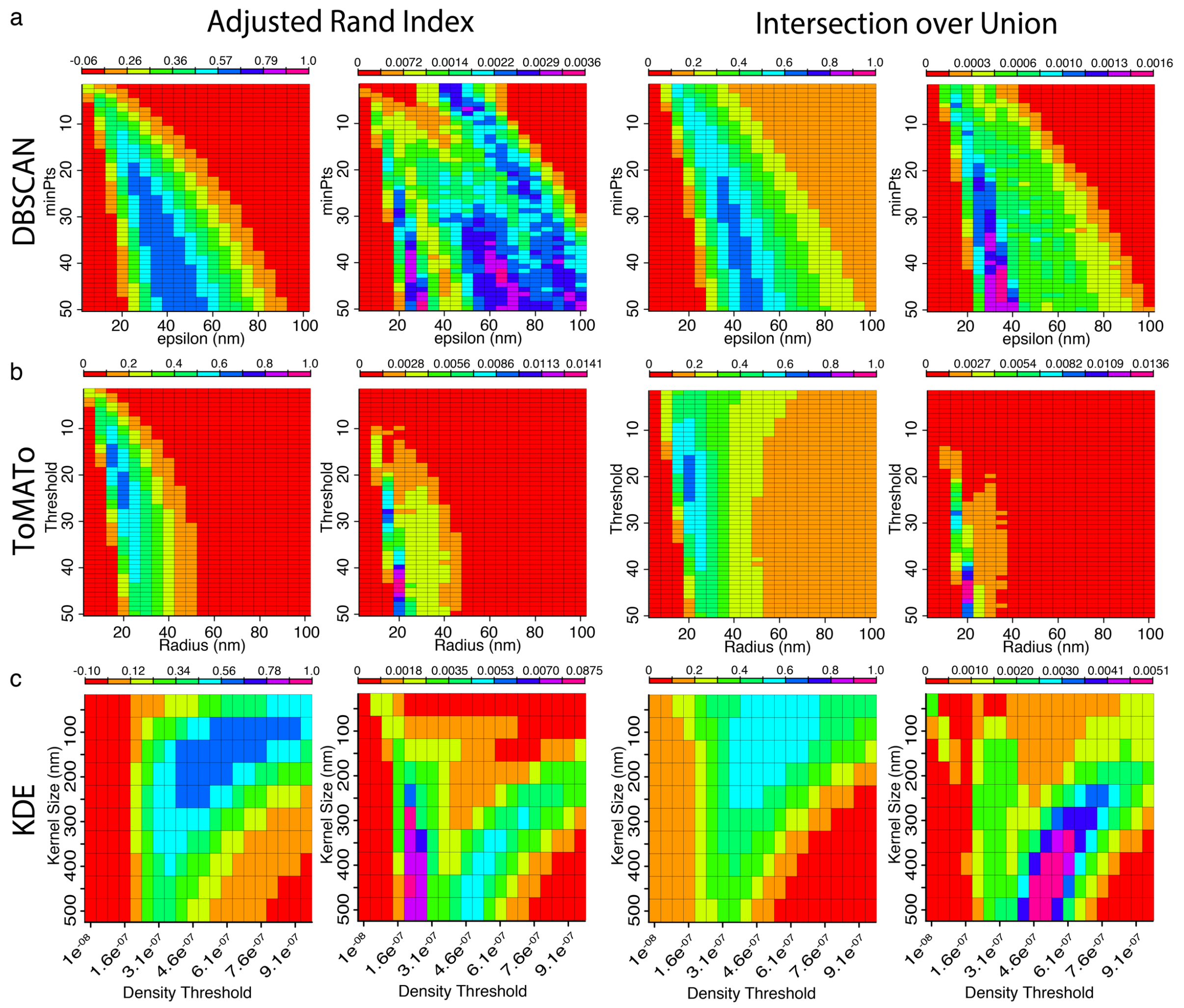


**Supplementary Figure 14. Parameter scan ARI and IoU matrices for DBSCAN, ToMATo, and KDE of elliptically shaped clusters, with fluorophore multiple blinking simulated (*Scenario 6*).** a) Parameter scan matrices for DBSCAN assessed by the ARI (left) and IoU (right). For each index we have the mean index value matrix, i.e., the mean index value from the 50 simulations for each parameter combination (left), and the variance matrix, which is the variance of the mean index matrix values (right). b) Parameter scan matrices for ToMATo assessed by the ARI (left) and IoU (right), with the mean index value matrix (left), and the variance matrix (right). c) Parameter scan matrices for KDE assessed by the ARI (left) and IoU (right), with the mean index value matrix (left), and the variance matrix (right).

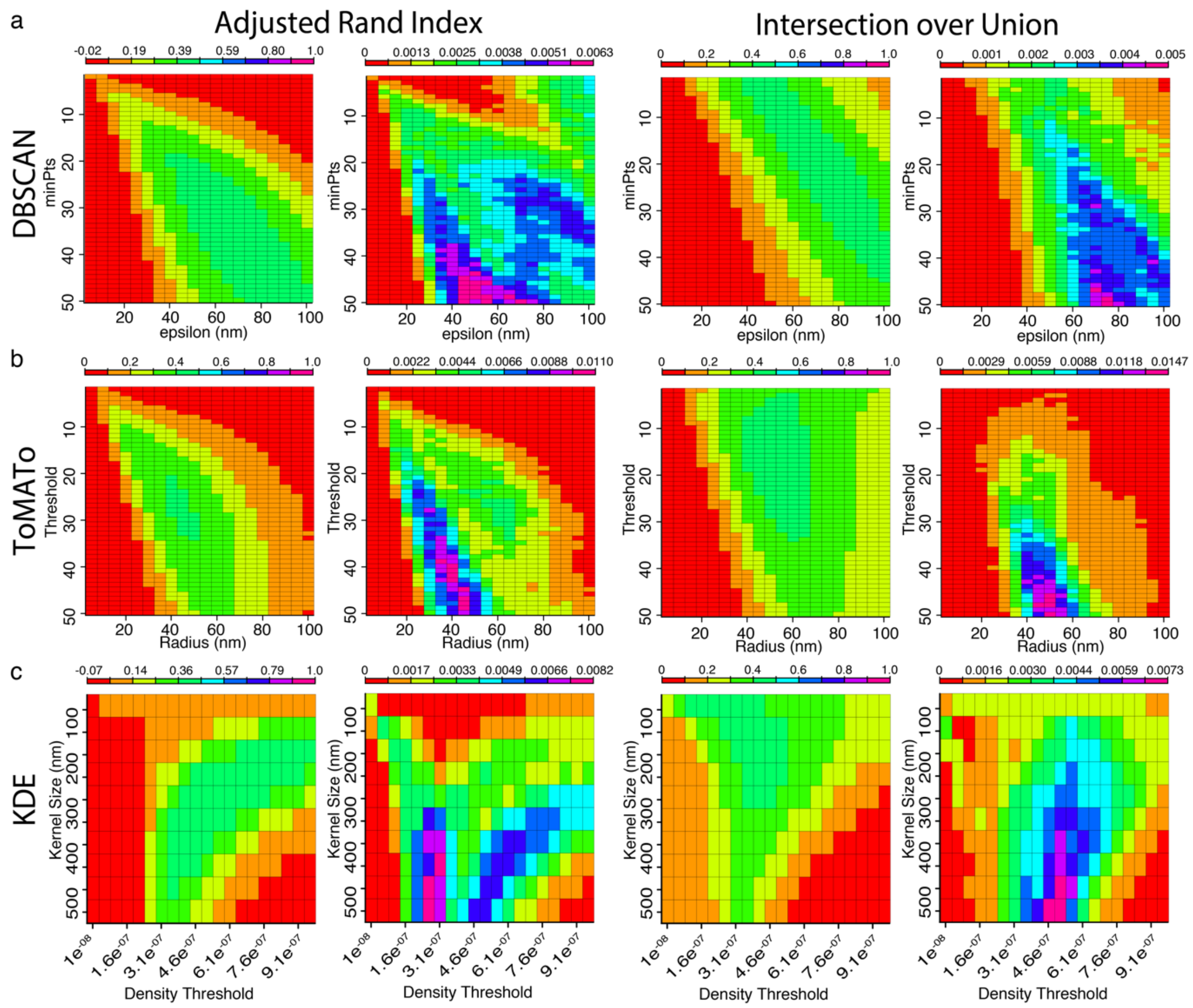


**Supplementary Figure 15. Parameter scan ARI and IoU matrices for DBSCAN, ToMATo, and KDE of clusters with different distribution widths, with fluorophore multiple blinking simulated (*Scenario 7*).** a) Parameter scan matrices for DBSCAN assessed by the ARI (left) and IoU (right). For each index we have the mean index value matrix, i.e., the mean index value from the 50 simulations for each parameter combination (left), and the variance matrix, which is the variance of the mean index matrix values (right). b) Parameter scan matrices for ToMATo assessed by the ARI (left) and IoU (right), with the mean index value matrix (left), and the variance matrix (right). c) Parameter scan matrices for KDE assessed by the ARI (left) and IoU (right), with the mean index value matrix (left), and the variance matrix (right).

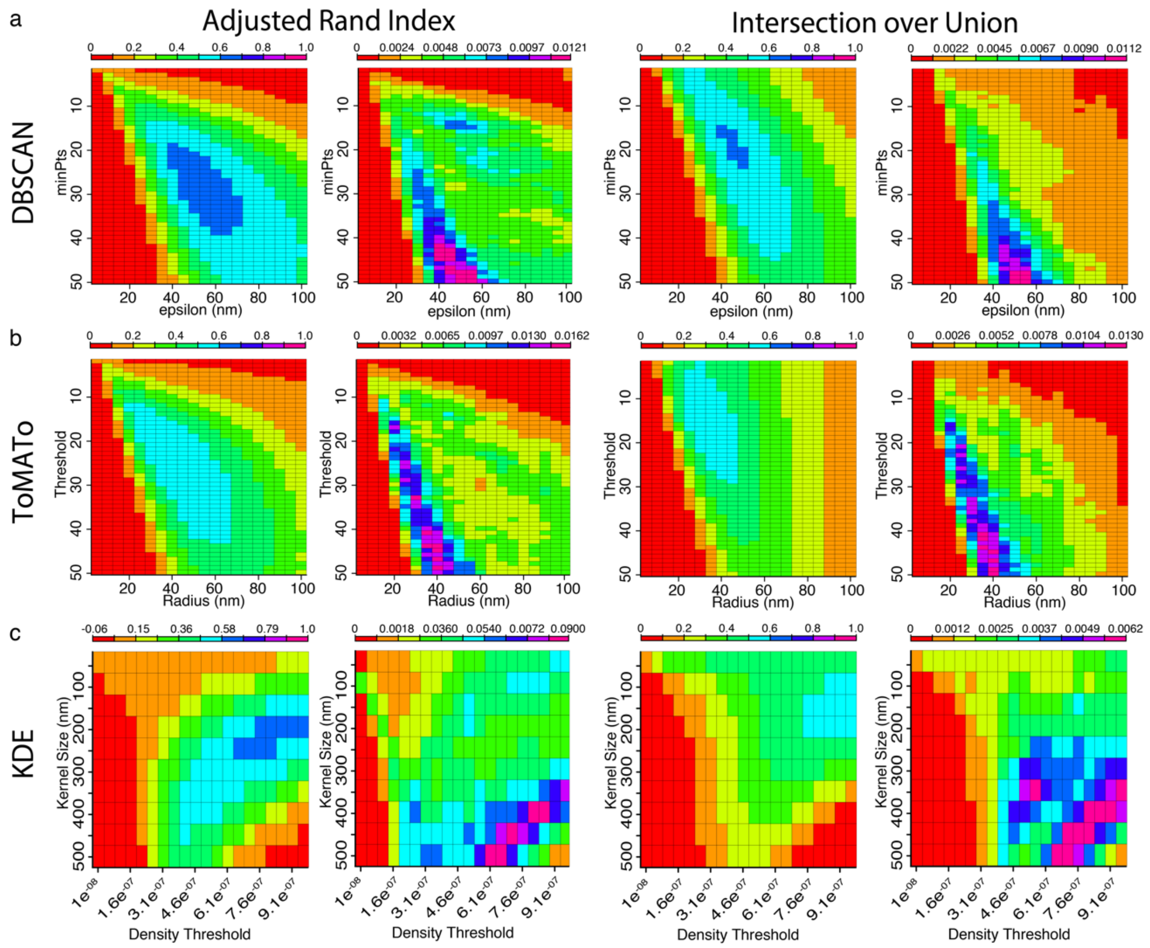


**Supplementary Figure 16. Parameter scan ARI and IoU matrices for DBSCAN, ToMATo, and KDE of clusters with different denisties, with fluorophore multiple blinking simulated (*Scenario 8*).** a) Parameter scan matrices for DBSCAN assessed by the ARI (left) and IoU (right). For each index we have the mean index value matrix, i.e., the mean index value from the 50 simulations for each parameter combination (left), and the variance matrix, which is the variance of the mean index matrix values (right). b) Parameter scan matrices for ToMATo assessed by the ARI (left) and IoU (right), with the mean index value matrix (left), and the variance matrix (right). c) Parameter scan matrices for KDE assessed by the ARI (left) and IoU (right), with the mean index value matrix (left), and the variance matrix (right).

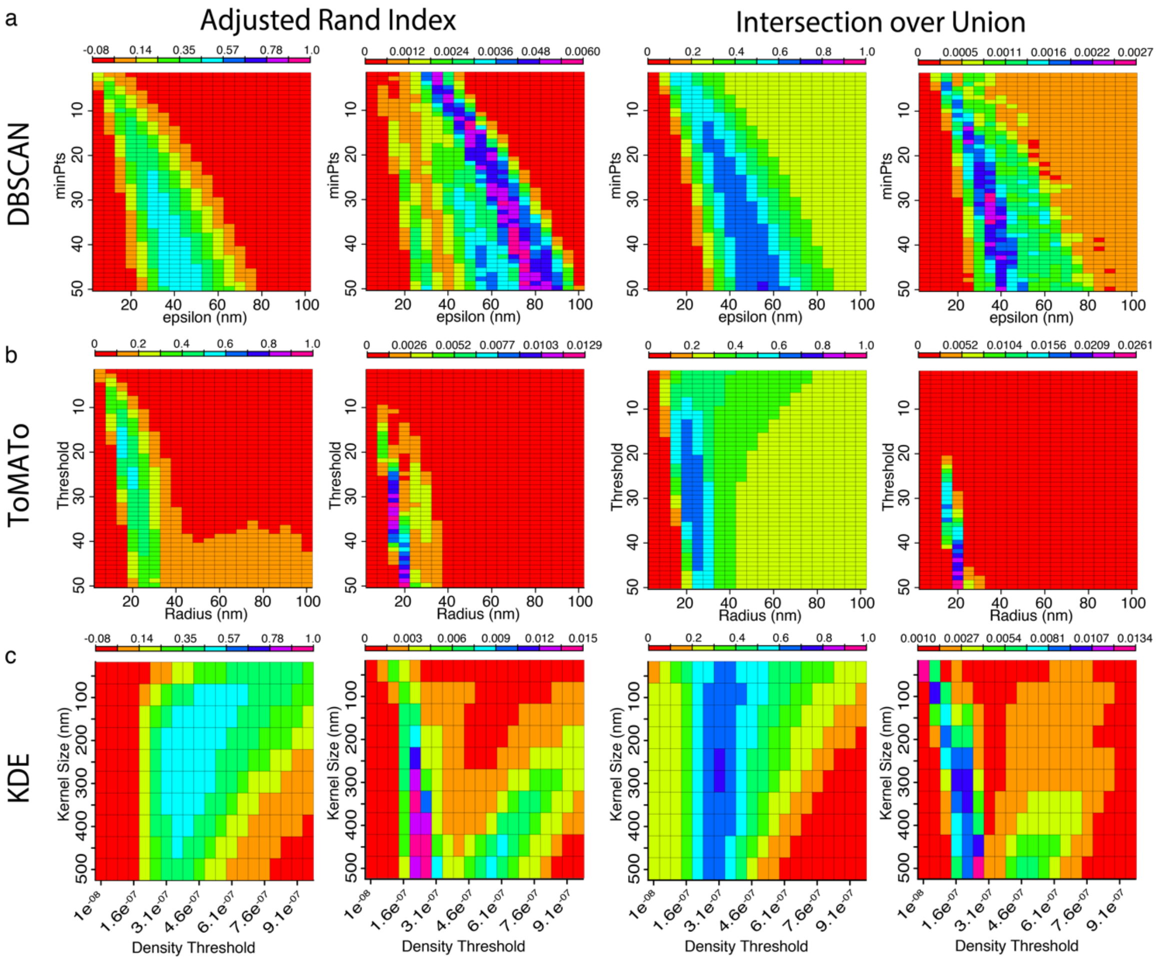


**Supplementary Figure 17. Parameter scan ARI and IoU matrices for DBSCAN, ToMATo, and KDE of clusters with different widths but density maintained, with fluorophore multiple blinking simulated (*Scenario 9*).** a) Parameter scan matrices for DBSCAN assessed by the ARI (left) and IoU (right). For each index we have the mean index value matrix, i.e., the mean index value from the 50 simulations for each parameter combination (left), and the variance matrix, which is the variance of the mean index matrix values (right). b) Parameter scan matrices for ToMATo assessed by the ARI (left) and IoU (right), with the mean index value matrix (left), and the variance matrix (right). c) Parameter scan matrices for KDE assessed by the ARI (left) and IoU (right), with the mean index value matrix (left), and the variance matrix (right).

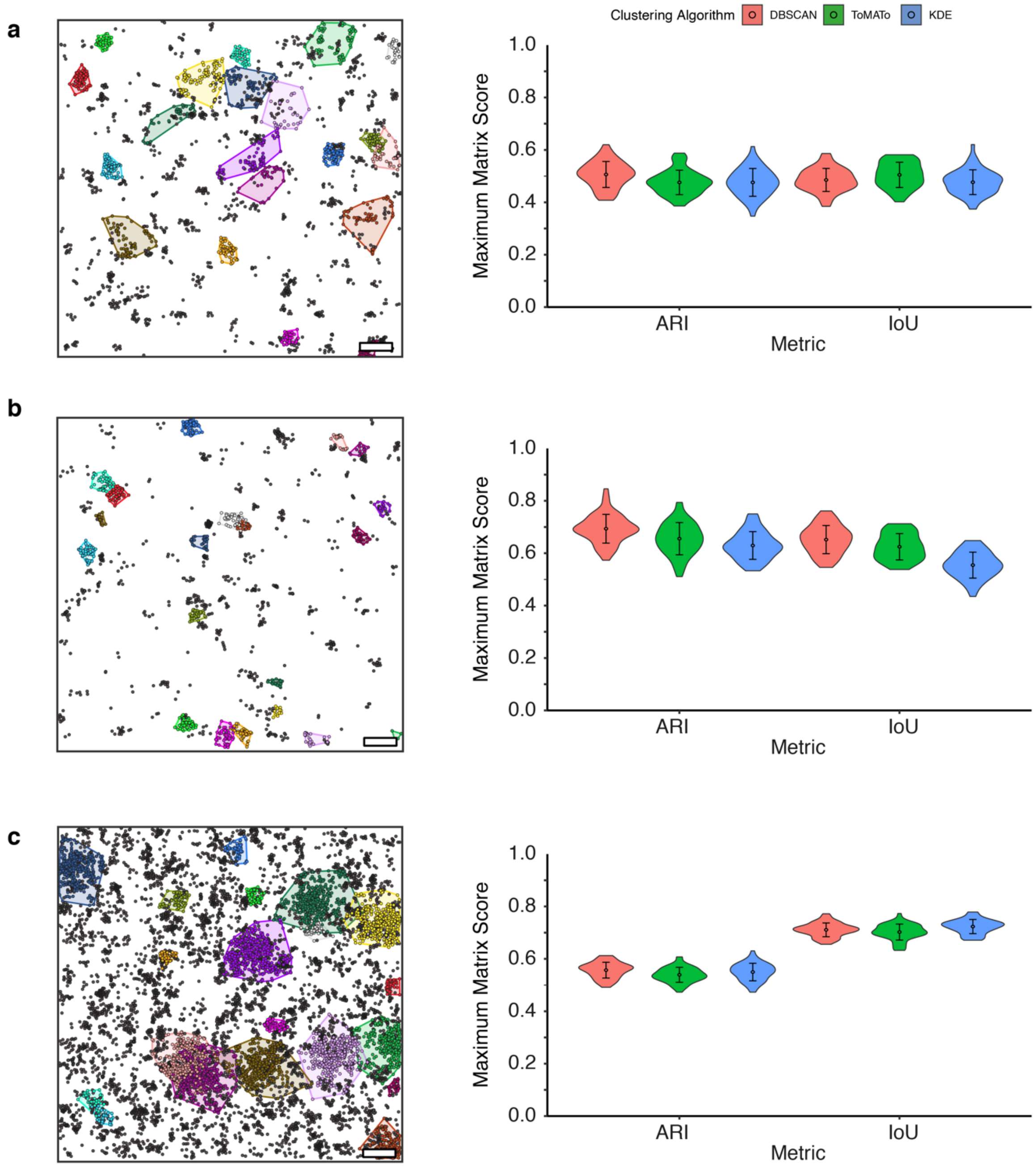


**Supplementary Figure 18. Performance of DBSCAN, ToMATo and KDE against Scenarios 7-9 in the presence of multiple fluorophore blinking.** Left: Representative ground truth clustering showing clustered points (multicolor with bounding polygons) and non-clustered localisations (black). Scale bar – 200 nm. Right: Maximum ARI and IoU scores. **a)** *Scenario 7*. **b)** *Scenario 8*. **c)** *Scenario 9*.

**Supplementary Table 1. Summary of the optimal clustering parameters for all algorithms in all clustered simulation scenarios.**

| | | | Clustering Algorithm | | | | | |
|---|---|---|---|---|---|---|---|---|
| | | | DBSCAN | | ToMATo | | KDE | |
| Simulation Scenario | Added blinking? | Index | minPts | epsilon (nm) | Threshold | Radius (nm) | Kernel size (nm) | Density Cut off |
| ***Scenario 2*** | No | ARI | 7 | 45 | 6 | 40 | 150 | $8.6e^{-7}$ |
| | | IoU | 7 | 45 | 6 | 40 | 100 | $9.6e^{-7}$ |
| | Yes | ARI | 35 | 50 | 20 | 30 | 200 | $7.6e^{-7}$ |
| | | IoU | 25 | 45 | 18 | 30 | 150 | $8.1e^{-7}$ |
| ***Scenario 3*** | No | ARI | 10 | 40 | 7 | 30 | 150 | $5.6e^{-7}$ |
| | | IoU | 10 | 40 | 7 | 30 | 100 | $6.1e^{-7}$ |
| | Yes | ARI | 50 | 45 | 14 | 15 | 150 | $6.1e^{-7}$ |
| | | IoU | 43 | 45 | 12 | 15 | 150 | $5.6e^{-7}$ |
| ***Scenario 4*** | No | ARI | 4 | 50 | 4 | 45 | 150 | $8.1e^{-7}$ |
| | | IoU | 5 | 55 | 4 | 50 | 100 | $9.6e^{-7}$ |
| | Yes | ARI | 25 | 60 | 26 | 50 | 250 | $6.1e^{-7}$ |
| | | IoU | 13 | 40 | 12 | 35 | 50 | $9.6e^{-7}$ |
| ***Scenario 5*** | No | ARI | 8 | 35 | 5 | 25 | 100 | $4.6e^{-7}$ |
| | | IoU | 8 | 35 | 5 | 30 | 50 | $5.1e^{-7}$ |
| | Yes | ARI | 34 | 35 | 13 | 15 | 100 | $4.6e^{-7}$ |
| | | IoU | 27 | 35 | 9 | 15 | 100 | $3.6e^{-7}$ |
| ***Scenario 6*** | No | ARI | 6 | 30 | 6 | 30 | 100 | $6.1e^{-7}$ |
| | | IoU | 7 | 35 | 6 | 30 | 50 | $7.1e^{-7}$ |
| | Yes | ARI | 42 | 40 | 16 | 15 | 150 | $5.6e^{-7}$ |
| | | IoU | 42 | 45 | 21 | 20 | 150 | $4.6e^{-7}$ |
| ***Scenario 7*** | No | ARI | 6 | 65 | 6 | 60 | 200 | $4.6e^{-7}$ |
| | | IoU | 6 | 80 | 6 | 70 | 150 | $4.1e^{-7}$ |
| | Yes | ARI | 32 | 65 | 29 | 45 | 100 | $5.6e^{-7}$ |
| | | IoU | 19 | 70 | 19 | 50 | 250 | $3.6e^{-7}$ |
| ***Scenario 8*** | No | ARI | 4 | 50 | 4 | 50 | 150 | $8.1e^{-7}$ |
| | | IoU | 4 | 45 | 4 | 45 | 500 | $9.1e^{-7}$ |
| | Yes | ARI | 27 | 50 | 26 | 30 | 200 | $9.6e^{-7}$ |
| | | IoU | 19 | 45 | 13 | 45 | 150 | $9.1e^{-7}$ |
| ***Scenario 9*** | No | ARI | 12 | 45 | 7 | 30 | 150 | $3.6e^{-7}$ |
| | | IoU | 45 | 95 | 15 | 40 | 250 | $2.6e^{-7}$ |
| | Yes | ARI | 50 | 45 | 18 | 15 | 200 | $4.1e^{-7}$ |
| | | IoU | 50 | 55 | 27 | 20 | 300 | $3.1e^{-7}$ |
